# Genome-scale requirements for dynein-based trafficking revealed by a high-content arrayed CRISPR screen

**DOI:** 10.1101/2023.03.01.530592

**Authors:** Chun Hao Wong, Steven W. Wingett, Chen Qian, J. Matthew Taliaferro, Douglas Ross-Thriepland, Simon L. Bullock

## Abstract

The cytoplasmic dynein-1 (dynein) motor plays a key role in cellular organisation by transporting a wide variety of cellular constituents towards the minus ends of microtubules. However, relatively little is known about how the biosynthesis, assembly and functional diversity of the motor is orchestrated. To address this issue, we have conducted an arrayed CRISPR loss-of-function screen in human cells using the distribution of dynein-tethered peroxisomes and early endosomes as readouts. From a guide RNA library targeting 18,253 genes, 195 validated hits were recovered and parsed into those impacting multiple dynein cargoes and those whose effects are restricted to a subset of cargoes. Clustering of high-dimensional phenotypic fingerprints generated from multiplexed images revealed co-functional genes involved in many cellular processes, including several candidate novel regulators of core dynein functions. Mechanistic analysis of one of these proteins, the RNA-binding protein SUGP1, provides evidence that it promotes cargo trafficking by sustaining functional expression of the dynein activator LIS1. Our dataset represents a rich source of new hypotheses for investigating microtubule-based transport, as well as several other aspects of cellular organisation that were captured by our high-content imaging.

## INTRODUCTION

Cytoskeletal motor proteins play a central role in organising the intracellular space. The cytoplasmic dynein-1 motor (dynein) is responsible for almost all motility towards the minus ends of microtubules and consequently carries a large variety of cargoes – including endosomes, lysosomes, autophagosomes, mitochondria, mRNAs and pathogens – towards the interior of the cell ^1^.

Dynein is a 1.3-MDa multi-subunit protein complex ^1, 2^. Its motor and microtubule-binding activities are housed in the C-terminal region of the heavy chain subunit, DYNC1H1. The N-terminal region of DYNC1H1 mediates self-dimerisation and provides a scaffold for docking of the accessory chains – two copies each of an intermediate chain (DYNC1I1 or DYNC1I2) and a light intermediate chain (DYNC1LI1 or DYNC1LI2), and six copies of a light chain (DYNLL1, DYNRB1 or DYNLT1). These subunits stabilise the heavy chain assembly and contribute to cargo recognition.

*In vitro* reconstitution experiments have shown that motility of human dynein is dependent on another large, multi-subunit complex – dynactin – and one of a number of coiled-coil-containing cargo adaptors (termed ‘activating adaptors’) ^1, 3-7^. The activating adaptors stabilise the interaction of dynein with dynactin, which switches on processive movement. Whilst some adaptors can interact with dynein and dynactin without additional factors ^8, 9^, others – such as the Golgi-vesicle-associated protein BICD2 – require an autoinhibitory state to be released by binding to a cargo-associated protein ^10-12^. Collectively, these mechanisms co-ordinate activity of the motor with the availability of cargo.

The Lissencephaly-1 (LIS1/PAFAH1B1) protein also plays a critical role in cargo transport. This factor binds the motor domain of DYNC1H1 and promotes formation of the dynein-dynactin-activating adaptor assembly ^13-19^. The importance of LIS1 is underlined by the finding that even modest reductions in its abundance impair dynein function and cause severe neurological disease ^20-23^.

Whilst *in vitro* studies have greatly advanced our understanding of dynein activation, many questions remain about how cargo trafficking by this motor is orchestrated in the cellular environment. For example, how is biosynthesis of individual components of the transport machinery, as well as their assembly into larger complexes, controlled? And how is the functional diversity of dynein achieved: are there mechanisms that regulate the behaviour of dynein complexes bound to specific cargoes?

To gain a foothold into these and other aspects of dynein biology, we have conducted a genome-wide loss-of-function CRISPR screen for factors that alter localisation of the motor’s cargoes in human cells and performed high-dimensional phenotypic analysis of screen hits. Our results represent a valuable resource for dissecting microtubule-based trafficking mechanisms, as well as several other aspects of cellular organisation that were captured in our images.

## RESULTS

### Optimised procedures for CRISPR/Cas9-mediated gene disruption in an arrayed format

We first sought to establish highly scalable procedures for CRISPR/Cas9-mediated gene editing in an arrayed format, *i*.*e*. in which one gene is targeted per well. Screening in this manner, as opposed to the more conventional pooled format, facilitates the use of complex, multivariate imaging readouts, as well as the establishment of phenotype-genotype relationships ^24^.

We developed a protocol in which a large pool of cells is transfected with mRNA encoding Cas9 and seeded into 384-well plates that have each well pre-dispensed with synthetic two-part guide RNAs (hereafter crRNAs) ^25, 26^ targeting a different gene (Figure 1A). To increase the frequency of gene disruption, four unique crRNAs were used per gene. Synthetic guide RNAs are well suited for arrayed screening due to their adaptability for plate layout manipulation. Delivering Cas9 by mRNA transfection circumvents the need to make stable cell lines expressing the enzyme from a genomic DNA construct, whilst generating a pool of mRNA-Cas9-expressing cells as starting material for crRNA delivery removes well-to-well differences in Cas9 transfection as a variable.

**Figure 1.**
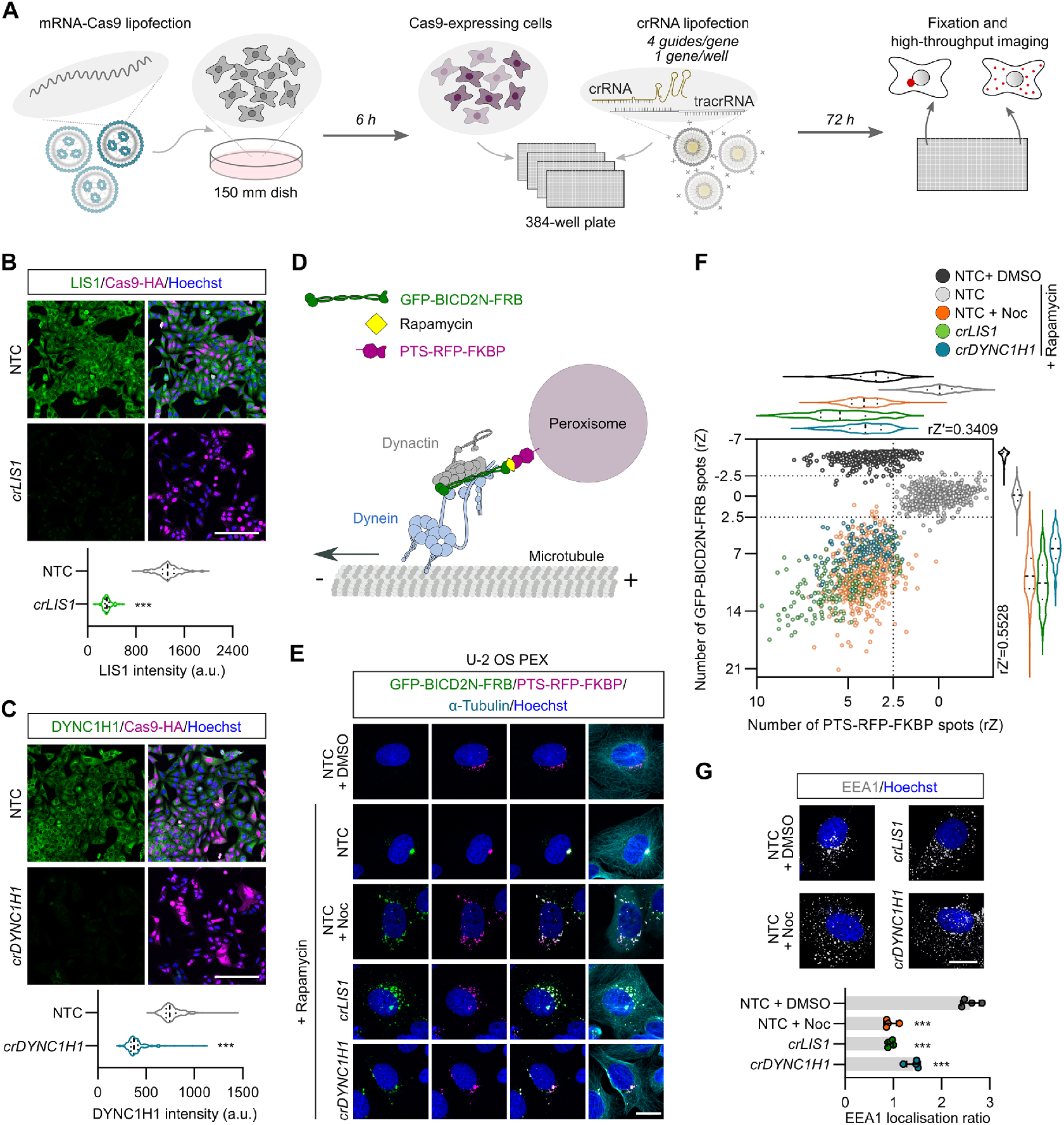
Assay development for arrayed CRISPR/Cas9 screening. **A)** Workflow for image-based screening of dynein cargo localisation. **B, C)** Representative images and quantification of immunostained unmodified U-2 OS cells following CRISPR/Cas9-mediated editing of *LIS1* (B) or *DYNC1H1* (C). NTC, non-targeting control; Hoescht, DNA stain; *cr, crRNA*. Violin plots show fluorescence intensity values at the single cell level (minimum of 100 cells from at least four wells for each group; median, bold line; first/third quartile, dashed lines). ***p<0.001 (two-tailed Mann-Whitney-test). Scale bar, 200 μm. **D)** Illustration of inducible peroxisome relocalisation assay. Only one motor complex is depicted per peroxisome for simplicity. **E)** Representative images of U-2 OS PEX cells stained for microtubules (α-Tubulin) and DNA (Hoechst) after the indicated treatments. Cells were treated with either DMSO (vehicle), rapamycin alone or rapamycin with nocodozole (Noc) for 2.5 h before fixation. Scale bar, 20 μm. **F)** Validation of inducible peroxisome relocalisation assay in high-throughput format. Scatter plot and corresponding violin plots (median, bold line; first/third quartile, dashed lines) of number of GFP-BICD2N-FRB and PTS-RFP-FKBP spots. Data points represent robust-Z (rZ) normalisation (central reference = NTC treated with rapamycin; value increases with cargo dispersion) with mean aggregation at well level (minimum of 100 wells analysed from 3 × 384-well plates). rZ′ values show assay window between NTC with rapamycin and *crLIS1* with rapamycin. **G)** Representative images and quantification of early endosome (EEA1) dispersion in unmodified U-2 OS cells after indicated treatments. Bar graph shows ratio between EEA1 spot number at the perinuclear region vs the peripheral region (lower values indicate increased dispersion). Data points represent mean aggregation at the well level (minimum of 100 cells analysed per well, four wells analysed per condition). Error bars, S.D.. ***p<0.001 (one-way ANOVA with Dunnett’s multiple comparison vs NTC + DMSO). Scale bar, 20 μm.

We optimised the mRNA-Cas9 transfection protocol in a panel of five commonly used human cell lines derived from different organs (U-2 OS, ARPE-19, HEK-293, IMR-90 and SH-SY5Y; Supplementary figure 1). These experiments defined mRNA concentrations, transfection reagents and transfection conditions that gave a very high proportion of Cas9-expressing cells (90-100%) yet had minimal toxicity. Editing efficiency with the optimised conditions was evaluated in ARPE-19 and U-2 OS cells using crRNAs targeting six genes, including *LIS1, DYNC1H1* and *DCTN1*, which encodes a dynactin subunit (*crLIS1, crDYNC1H1* and *crDCTN1*, respectively). In ARPE-19 cells, 70-80% of cells had strongly reduced expression of the protein products of the target genes 72 hours after crRNA transfection (Supplementary figure 2A), whereas in U-2 OS cells this value was 85-100% (Figure 1B, C and Supplementary figure 2B). Thus, our RNA-mediated reagent delivery methods disrupt a range of target genes with high efficiency. These experiments also demonstrated that a 72-hour targeting window allows retention of sufficient edited cells when targeting essential genes that disrupt dynein-based transport.

### Imaging-based assays for dynein activity

As the above experiments revealed particularly high rates of gene editing in U-2 OS cells, we sought to develop an imaging-based readout of dynein activity in this system that would be suitable for an arrayed screen. We took advantage of a previously characterised U-2 OS line (hereafter U-2 OS PEX) ^27^ that has a chemically-inducible system for dynein-mediated relocalisation of fluorescent peroxisomes ^12, 27, 28^. This line stably expresses the constitutively active N-terminal region of the activating adaptor BICD2 (BICD2N) fused to GFP and FRB (GFP-BICD2N-FRB), as well as a peroxisome targeting sequence (PTS) fused to RFP and FKBP (PTS-RFP-FKBP) (Figure 1D). Addition of rapamycin triggers association of BICD2N with peroxisomes via FRB-FKBP heterodimerisation ^29^, which recruits dynein and dynactin complexes to the organelle (Figure 1D). This leads to strong clustering of peroxisomes – which otherwise are dispersed in the perinuclear region – at the juxtanuclear microtubule-organising centre (MTOC), where microtubule minus ends are enriched (Figure 1E) ^12, 27, 28^.

The assay was optimised by automatically quantifying the number of GFP and RFP spots, which acts as a proxy for peroxisome clustering, in response to rapamycin concentration and incubation time, as well as the number of seeded cells (Supplementary figure 3). We also corroborated the previous observation that rapamycin-induced relocalisation of peroxisomes in U-2 OS PEX cells is impaired by depolymerisation of microtubules with nocodazole ^27^ and demonstrated this is also the case when either *LIS1* or *DYNC1H1* are targeted with our optimised CRISPR protocol (Figure 1E).

The assay was subsequently scaled and validated through a ‘Min-Max’ analysis in the presence of rapamycin with multiple plates pre-dispensed with rows of crRNAs targeting *LIS1, DYNC1H1* or *PLK1*(disruption of which blocks cell proliferation and thus serves as a label-free control for editing efficiency ^30, 31^). As controls, wells were dispensed with crRNAs that lack targets in the human genome (non-targeting control; NTC), or treated with nocodazole. Highly efficient gene disruption was observed across the plates for all three gene targets (Supplementary figure 4). Furthermore, there was a consistent change in the number of RFP and GFP spots in *crLIS1, crDYNC1H1* and nocodozole-treated wells compared to NTC (Figure 1F), demonstrating robust dispersal phenotypes. The assay window measured by the robust Z-prime (rZ′) between NTC and *crLIS1* was 0.34 and 0.55 for the RFP and GFP data, respectively, indicating suitability for imaging-based screening ^32^.

To maximise the information gained from a genome-wide CRISPR screen, we sought to monitor localisation of a second dynein cargo. We selected early endosomes, which rely on both dynein and dynactin for enrichment in the perinuclear region of the cytoplasm ^33-35^. Antibody staining against early endosome antigen 1 (EEA1) protein confirmed that nocodazole treatment, as well as disruption of *DYNC1H1* or *LIS1* by CRISPR, disperses early endosomes in U-2 OS cells, and this effect was quantified by measuring the ratio of EEA1 spots in the perinuclear versus the peripheral region of the cytoplasm (Figure 1G). Unlike the fluorescent peroxisomes in the U-2 OS PEX line, dynein-dynactin is linked to early endosomes by activating adaptors of the HOOK family ^36-39^. Thus, simultaneously screening for defects in peroxisome and early endosome localisation can potentially reveal factors involved in trafficking by discrete dynein-dynactin-activating adaptor complexes.

### Execution of the genome-wide screen and performance assessment

For the genome-wide screen, we adapted the aforementioned assays for peroxisome and early endosome distribution for end-to-end execution via automated liquid handlers. Screening was performed across 61 unique 384-well plates that were arrayed with a commercial crRNA library targeting 18,253 genes (four guides per gene). Neutral (NTC) and positive (*crLIS1* and *crPLK1*) controls were included in each plate for downstream quality assessment and normalisation (Figure 2A).

**Figure 2.**
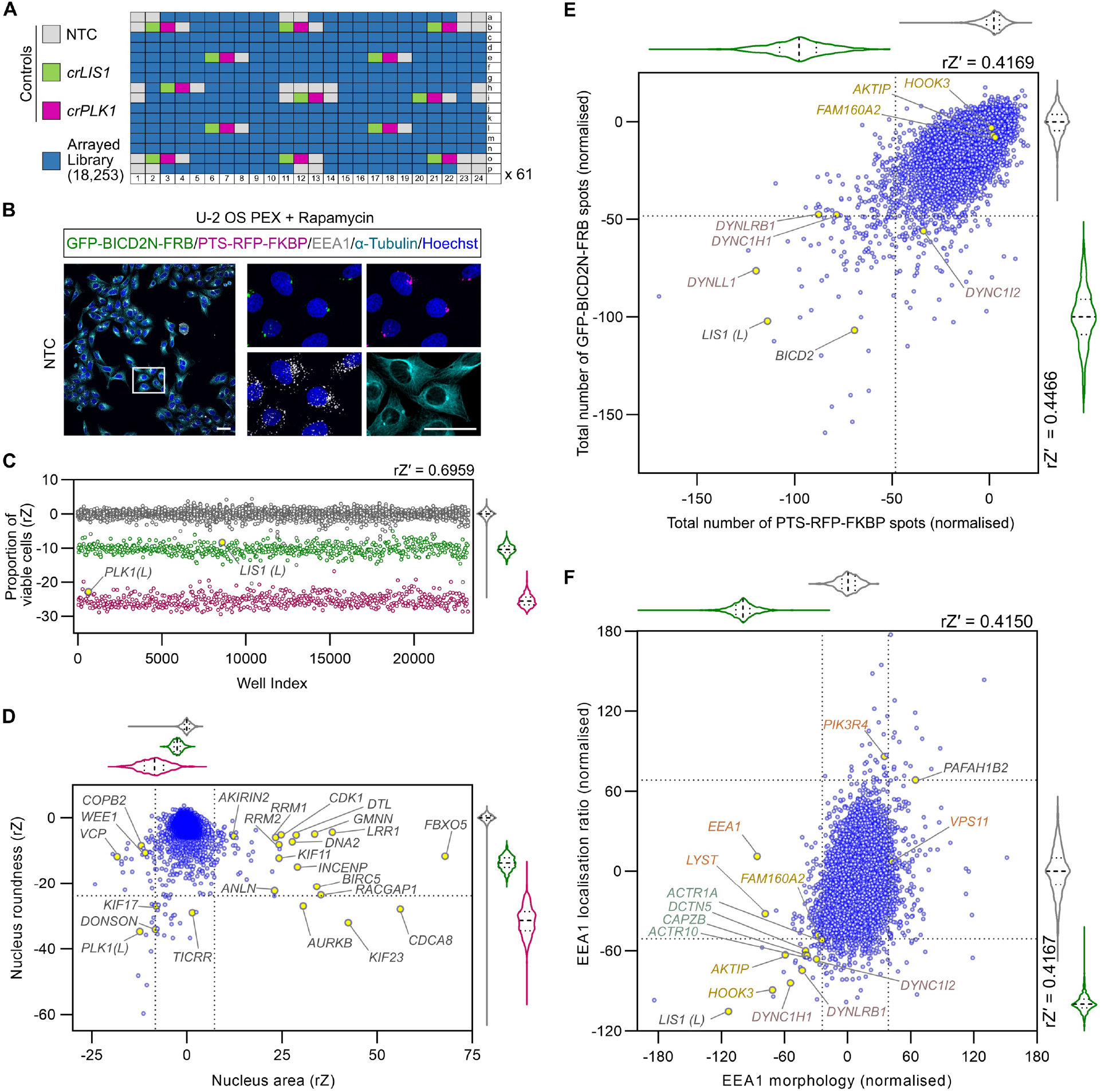
The genome-wide screen recovers known components of the dynein-dynactin machinery, as well as novel hits. **A)** Plate layout for genome-wide screen. **B)** Example of imaging data from the screen (maximum intensity projections of Z-stacks captured using a 20x/1.0 NA water objective). Scale bar, 50 μm. **C)** Evaluation of editing efficiency across the screen using cell viability as a readout. Scatter plot and corresponding violin plot for controls (median, bold dashed line; first/third quartile, dashed lines) of proportion of viable cells (gated based on nuclear morphology of NTC cells) per well after treatment with NTC, *crLIS1* or *crPLK1*. Data points (colour coded as in panel A) are rZ normalised values (central reference = NTC). rZ′ value shows assay window between NTC and *crPLK1*. The library copies of *crPLK1* and *crLIS1* are labelled with (L). **D)** Effects of arrayed library crRNAs on area and roundness of nuclei. Scatter plot and corresponding violin plots (median, bold dashed line; first/third quartile, dashed lines) of rZ normalised values (colour coded as in A; central reference = NTC). Dashed lines represent ± 4*S.D. of NTC and *crLIS1* (x-axis) or - 3.5*S.D. of NTC and *crLIS1* (y-axis), which were thresholds for hit calling. Genes previously shown to influence nuclear morphology are highlighted. **E, F)** Example of endpoints used for hit selection from peroxisome (E) and early endosome (F) data. Scatter plot and corresponding violin plots (median, bold dashed line; first/third quartile, dashed lines) of normalised values (colour coded as in A) based on the neutral control (NTC, 0) and inhibition (*crLIS1*, -100); rZ′ values show assay window between NTC and *crLIS1* control wells. Dashed lines on the x- and y-axes represent, respectively, - 2*S.D. (E) or - 3*S.D. (F) thresholds for hit calling. In F, ‘EEA1 morphology’ values were generated from a linear discriminant analysis weighted average of three endpoints quantifying the symmetry, intensity profile and texture of the EEA1 signal. ‘EEA1 localisation ratio’ is the ratio between the number of EEA1 spots at the perinuclear region vs at the peripheral region. Core components of the dynein complex and dynactin complex that met the threshold for hit calling are labelled in purple and teal text, respectively. Other categories of genes highlighted in the text are labelled in gold or orange. See Supplementary table 1 for source data for C-F.

The assay involved imaging mRNA-Cas9-transfected U-2 OS PEX cells that were fixed 72 hours after crRNA transfection and 2.5 hours after rapamycin addition. Cells were then stained with antibodies to EEA1, along with antibodies to α-Tubulin and Hoechst for cell segmentation. The resulting signals, together with those from the GFP and RFP channels, were captured with a high-content imaging platform (Figure 2B). In total, 8,150,065 cells from 24,576 wells (four fields of view per well; median of 345 cells analysed per well) were segmented for multi-parametric analyses. The complete set of quantitative data from the screen and the quality assessment of individual features used for hit calling are provided in Supplementary table 1 and Supplementary figure 5, respectively.

To gauge editing efficiency, we first evaluated the effects of the *crPLK1* controls on cell survival. As cell number is variable in the context of microplate-based assays, we performed population gating for viable cells based on Hoescht staining of DNA (*i*.*e*. removing any cells with apoptotic or mitotic features, or abnormal nuclear morphology (rZ′ = 0.7 (NTC versus *crPLK1*)). There was a large decrease in viable cells in *crPLK1* wells across the plates, as well as in the single library copy of *crPLK1* from the arrayed library (Figure 2C). In-keeping with a role of LIS1 in promoting dynein function in mitosis ^40, 41^, *crLIS1* controls across the plates, as well as the single library copy of *crLIS1*, also reduced the proportion of viable cells but to a lesser extent than *crPLK1* (Figure 2C). Together, these results indicate a high level of consistency in Cas9/crRNA activity across the screen.

In addition to *crPLK1*, crRNAs targeting 62 genes reduced cell viability to a significantly greater extent that *crLIS1*. Approximately 75% of these genes were previously found to be essential in cancer cell lines (Supplementary figure 6A) ^42^. Evaluating nuclear area and roundness across the complete set of assay plates identified many genes that have been implicated in the regulation of nuclear size in previous studies (Figure 2D) ^30, 43-45^. We also analysed the induction of micronuclei (Supplementary figure 6B), which to our knowledge has not been assessed in earlier screens. Many of the hits from this analysis encode essential components of the mitotic machinery, consistent with the contribution of chromosome segregation defects to micronuclei formation ^46^. In addition to recovering genes that were expected to influence cell survival, nuclear morphology and micronuclei formation, these analyses also implicated many other genes in these processes (Supplementary tables 1 and 2). Collectively, these observations show that our procedures effectively identify known, as well as novel, genotype-phenotype associations.

### Recovery of known and candidate novel players in dynein biology

To identify genes that are candidates to contribute to dynein-based trafficking, we performed multi-parametric analysis on the PTS-RFP-FBKP, GFP-BICD2N-FRB and EEA1 signals across the screening plates. crRNAs with strong effects on cell viability and nuclear morphology were excluded from further analysis, as we reasoned they may affect cargo localisation indirectly. Figure 2E shows the readout for the total number of GFP-BICD2N-FRB and PTS-RFP-FKBP spots per cell, which were amongst the metrics that gave a robust readout of peroxisome dispersion (rZ′ score of 0.41-0.45 comparing NTC and *crLIS1*). Genes encoding several components of the dynein complex, as well as *LIS1*, were identified amongst 217 factors whose targeting with library crRNAs caused a significant change in the number of either GFP and/or RFP spots (≥ ±2 S.D. of NTC). *BICD2* was also recovered as a hit in this analysis (Figure 2E),although this may reflect the ability of two of the crRNAs in this pool to recognise the GFP-BICD2N-FRB construct. We used several other metrics to quantify the subcellular localisation of GFP or RFP spots by segmenting the cytoplasmic region into perinuclear, intermediate and outer regions. This analysis recovered genes encoding other dynein and dynactin constituents (Supplementary table 3), as well as 114 additional genes not identified by the analysis of total spot number (Supplementary table 2).

Guides targeting dynein-dynactin components and *LIS1* were also part of 35 crRNA pools that significantly reduced perinuclear enrichment of EEA1-positive early endosomes (Figure 2F). The hits also included crRNAs to *HOOK3* and *AKTIP* (also known as *FTS*), which encode two of the proteins that form an ‘FHF’ complex linking dynein to early endosomes ^38^. The gene encoding the third complex component, *FAM160A2* (also known as *FHIP1B*), did not meet the threshold for inclusion as a hit for endosome dispersion but was very close to doing so. crRNAs for *HOOK3, AKTIP* and *FAM160A2* did not significantly affect peroxisome distribution (Figure 2E), consistent with the notion that their protein products function specifically in trafficking of early endosomes ^38^. We also found that several crRNA pools were associated with excessive clustering of early endosomes in the perinuclear region (Figure 2F), raising the possibility that their targets inhibit transport of early endosomes by dynein. These genes included *PAFAH1B2*, which encodes a catalytic subunit of the platelet-activating factor acetylhydrolase Ib complex that interacts directly with LIS1 ^47^. Our finding that *PAFAH1B2* disruption increases early endosome clustering supports the hypothesis that competition between PAFAH1B2 and dynein for LIS1 binding modulates motor activity ^48^. More generally, the identification of multiple known players in dynein-based transport demonstrates that our screening and analysis pipeline effectively identifies genes important for this process.

Our analysis additionally revealed crRNA pools that affected the morphology of peroxisomes (Supplementary figure 6C) and endosomes (Figure 2F) without changing their spatial distribution in the cell (Supplementary table 2). Some of these genes have a well-established link with peroxisome or endosome biology, notably the *PEX* genes and *DNML1*, which function in peroxisome biogenesis and fission, respectively ^49, 50^, and *VPS11, PIK3R4*, and *LYST*, which have roles in endosome biogenesis and/or endocytosis ^51, 52^. However, several other genes in this category have not previously been linked to peroxisome or endosome biology. Future studies of these genes could give new insight into the formation or turnover of these structures.

### Hit validation and coarse-grain phenotypic analysis

As our main objective was to identify factors important for dynein-based trafficking, we focused our downstream efforts on the screen hits that affected subcellular distribution of peroxisomes and/or early endosomes. We took forward a total of 376 hits in this category for validation in a secondary screen; 322 of these met our criteria for one or more metrics of peroxisome dispersion, 45 increased or decreased early endosome clustering and nine affected localisation of both cargoes (Supplementary table 4). We also took *FAM160A2* into the secondary screen because, as described above, this gene was very close to the hit threshold for EEA1 and has an established role in early endosome transport.

We retested the activity of the shortlisted crRNAs towards BICD2N-tethered peroxisomes in rapamycin-treated U-2 OS PEX cells, as well as towards early endosomes in an untreated, unmodified U-2 OS cell line. As the primary screen indicated specificity of some factors for a subset of cargoes, we also assessed the effects of each of the crRNA pools on perinuclear localisation of the Golgi apparatus (marked with TGN46 antibodies) in the untreated, unmodified cells. Studies in other cell types have shown that impairing dynein function causes dispersion of the Golgi ^53, 54^, and we confirmed this is also the case in U-2 OS cells using *crDYNC1H1* or *crLIS1* (Supplementary figure 7).

Each cargo was assayed in two independent screens, in which there was good agreement in general between the effects of the crRNA pools (Supplementary figure 8). The cut-offs applied previously to the genome-wide data were relatively lenient to maximise chances of capturing relevant hits in a ‘one-shot’ screening format. Here, we used more stringent gating, which led to the removal of 81 crRNAs that impacted cell viability and morphology based on the range observed with *crLIS1*. Of the remaining 296 crRNA pools, 195 caused a significant change in distribution of at least one of the cargoes (ratio of perinuclear to peripheral signal ±2.5 S.D. of NTC; Supplementary table 5). This validation rate (66%) is comparable to that of a previous arrayed CRISPR screen that used a subset of the same crRNA library to analyse delivery of lipid nanoparticle-encapsulated mRNA (50%) ^31^.

To better visualise the effects of the crRNAs on cargo distribution in the secondary screen, we used K-means clustering to group the 296 crRNA pools based on their mean effect sizes on localisation of peroxisomes, early endosomes and the Golgi (Figure 3). crRNAs targeting *AKTIP, FAM160A2* and *HOOK3* were found in the same cluster due to selective inhibition of early endosome clustering, whereas *crPAFAH1B2* was unique in strongly promoting clustering of early endosomes. The observation that targeting *PAFAH1B2* did not increase perinuclear localisation of peroxisomes and the Golgi may reflect these cargoes already being tightly clustered at this site in the wild-type situation. We also identified clusters of crRNAs that affected distribution of all three cargoes (Figure 3). These included cluster 14, which was associated with strong dispersion phenotypes and comprised crRNAs targeting *LIS1* and three dynein components, as well as five other genes, and cluster 13, which had more modest cargo dispersion and comprised four dynactin components and 16 other genes (see Supplementary table 5 for list of genes in each cluster). Other clusters contained crRNAs with more selective effects. These included cluster 9, which was characterised by dispersion of peroxisomes and the Golgi but not early endosomes. We conclude from this series of experiments that the genome-wide screen successfully identified genes that influence the distribution of dynein cargoes, including many that selectively affect a subset of cargo types.

**Figure 3.**
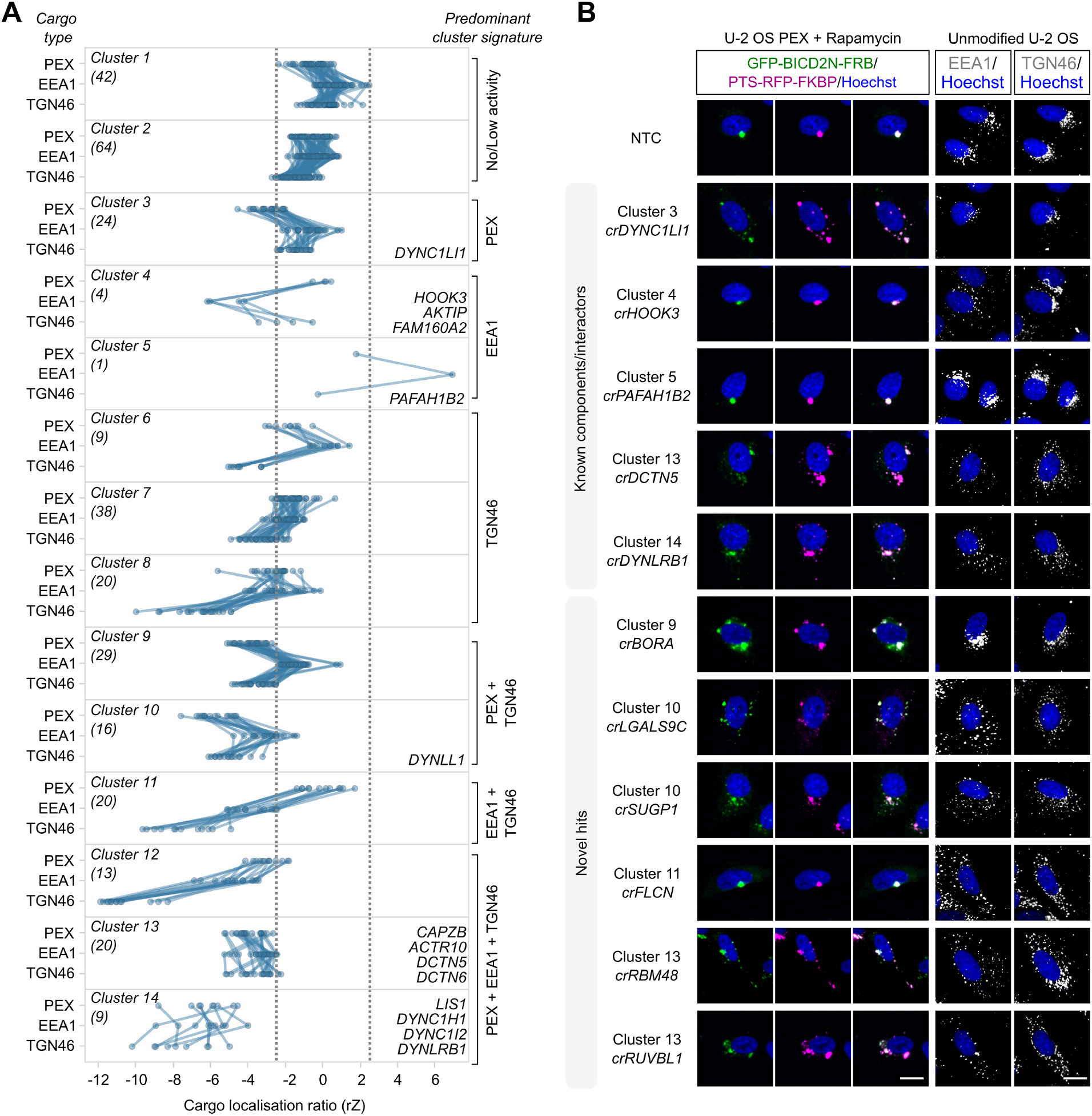
Screen hits can be categorised based on effects on different dynein cargoes. **A)** Grouping of hits based on effects in the secondary screen on localisation of peroxisomes (PEX; average of GFP-BICD2N-FRB and PTS-RFP-FKBP), early endosomes (EEA1) and *trans*-Golgi network (TGN46). Cargo localisation ratio was calculated by dividing spot number at the perinuclear region vs the peripheral region (negative values indicate increased dispersion). Grouping was performed with K-means clustering with Euclidean distance. Data points represent the average rZ (central reference = NTC) from two independent experiments (at least four wells per crRNA per experiment), with each line representing a crRNA pool. Dashed lines show 2.5*S.D.. Individual clusters are labelled with the number of constituent genes (parentheses), examples of constituent genes, and manual annotation of the predominant cargo signature. See Supplementary table 5 for list of genes in each cluster. **B)** Representative images of cargo localisation in cells edited with crRNAs targeting known components and well-characterised interactors of the dynein machinery, as well as novel hits. Third column from left is merge of GFP-BICD2N-FRB and PTS-RFP-FKBP signals. Scale bar, 20 μm.

To further evaluate the phenotypes in the secondary screen, we determined effects of the original shortlist of 377 crRNAs pools on the microtubule cytoskeleton by staining assay plates with γ-Tubulin or α-Tubulin antibodies. These reagents mark the MTOC and microtubule network, respectively. γ-Tubulin staining revealed that, compared to the NTC condition, 243 cRNA pools (64.5%) decreased the proportion of cells with one MTOC. Of these pools, 193 (51.2% of the total) increased the proportion of cells with more than one MTOC and 50 (13.3% of the total) increased the proportion with no MTOCs (Supplementary figure 9A and B; Supplementary table 6). Several of the pools that increased MTOC number targeted factors with well-established functions in mitosis, including PLK1, AURKA and CHMP4B ^55, 56^. The crRNAs that impaired MTOC formation included those targeting the centriole component SAS6 (*crSASS6*), the γ-tubulin ring complex (γ-TuRC) component GCP4 (*crTUBGCP4*), and subunits of the tubulin chaperone complex TRiC/CCT ^57-59^. crRNAs that target LIS1, the dynein subunits DYNC1H1, DYNC1I2, and the dynactin subunit ACTR1A also lowered MTOC number, in keeping with established roles of the motor complex in delivering structural and regulatory components of the centrosome ^60, 61^.

Staining the edited cells with α-Tubulin antibodies showed that none of the crRNAs caused a strong depletion of microtubules, such as that seen in nocodazole controls (Supplementary figure 9C; Supplementary table 6). However, 40 (11%) of the hits were associated with more subtle alterations in morphology of the microtubule network. These factors included genes encoding several tubulin isotypes ^62^ and TRiC/CCT components, as well as LIS1, the dynein light chain DYNLL1 and the dynactin component DCTN5. The observation of α-Tubulin phenotypes with crRNAs that target components of the dynein machinery is consistent with the motor’s ability to modulate interphase microtubule networks ^63-65^.

We conclude from our analysis of γ-Tubulin and α-Tubulin localisation that a subset of crRNA pools influence the microtubule cytoskeleton, including several that target core components of the dynein-dynactin complex.

### Identification of co-functional genes by unsupervised phenotypic clustering

The above analysis used a small number of features to give a coarse-grain phenotypic assessment of the hits. Our findings indicated that whereas core dynein-dynactin components affect the localisation of multiple cargoes and the organisation of the microtubule cytoskeleton, other hits from the screen participate in a subset of these processes. To systematically classify the function of the novel hits, we adapted an established image-based profiling workflow ^66-68^ to generate detailed phenotypic fingerprints for each crRNA pool. The fingerprints were then compared with each other to identify sets of genes that have similar phenotypes and which are therefore candidates to function in the same process.

To generate phenotypic fingerprints, we first extracted 2003 quantitative phenotypic parameters related to fluorescent signals from GFP-BICD2N-FRB, PTS-RFP-FKBP, EEA1, TGN46, α-Tubulin, γ-Tubulin and Hoescht (Figure 4A). The number of features was subsequently reduced to 278 by removing those that are highly variable or redundant via linear regression. Visualising the relative distribution of the high-dimensional phenotypic points on a two-dimensional Uniform Manifold Approximation and Projection (UMAP) plot revealed grouping of genes encoding members of the same protein complexes, such as histones, ribosomal proteins, RNA polymerase II, the RUVBL and TRiC/CCT chaperonins, FAM160A2-AKTIP-HOOK3 and dynein-dynactin (Figure 4B). Remarkably, there was also separate groupings of components of the core (A/B subunits) and regulatory particles (C/D subunits) of the proteasome ^69^. These observations highlight the utility of our procedures for identifying co-functional genes.

**Figure 4.**
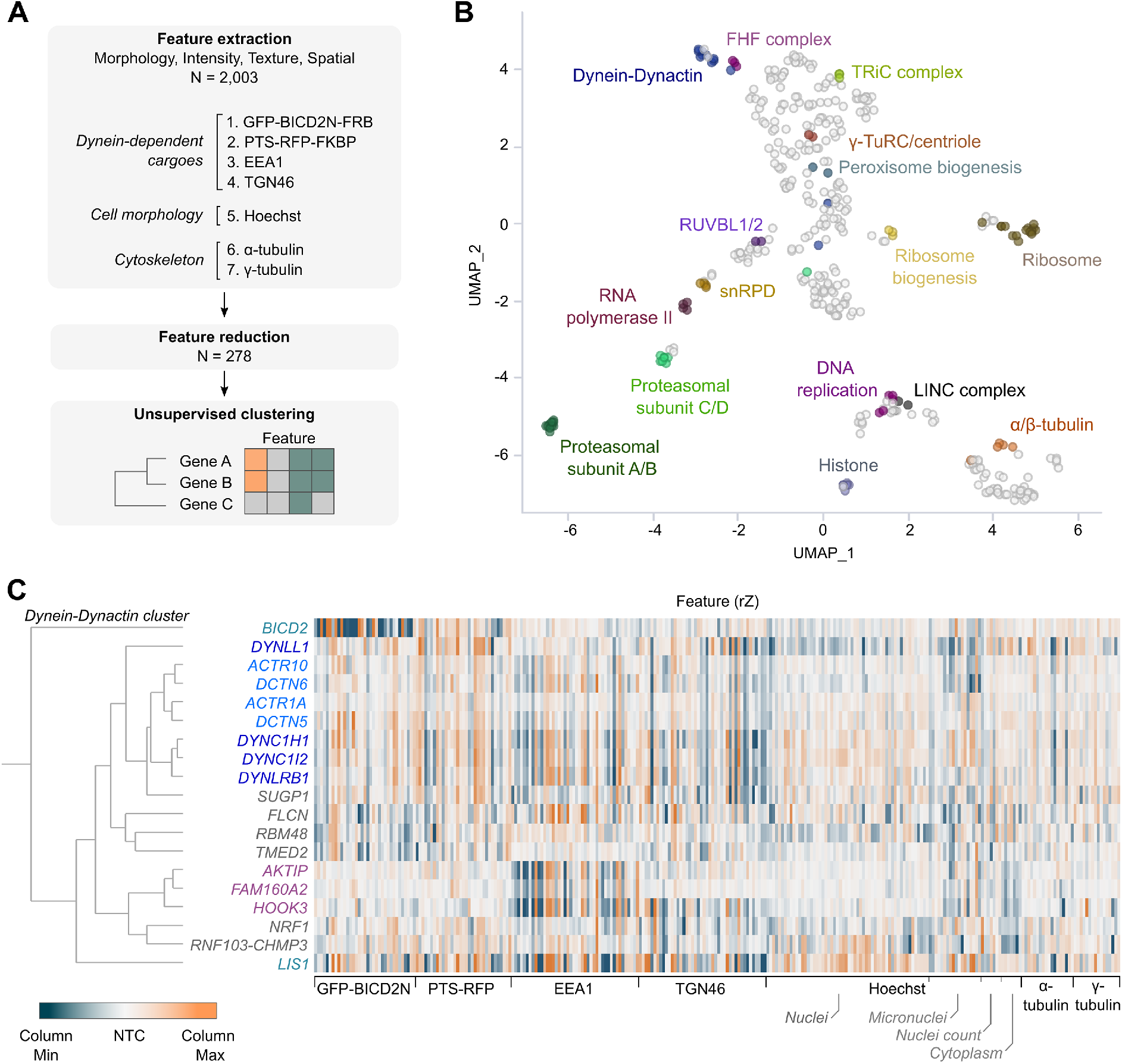
Unsupervised image-based profiling identifies a functional cluster containing known components of the dynein machinery and novel factors. **A)** Workflow for phenotypic profiling using images collected from the secondary screen. **B)** UMAP plot for phenotypes of genes selected from the primary screen. Clusters of genes with well-defined functions are highlighted. **C)** Phenotypic feature heatmap of the ‘dynein-dynactin’ gene cluster. Features are grouped in the x-axis according to the marker (see zoomable ‘Supplementary phenotypic heatmap’ file and Supplementary table 7 for names of individual features). Genes encoding dynein and dynactin components, as well as the associated proteins BICD2 and LIS1, are labelled in different shades of blue, whereas FHF component genes are shown in magenta. Novel genes in the cluster are labelled in grey. The scales of rZ values (central reference = NTC) were adjusted based on min and max values of individual features. ‘Cytoplasm’ refers to features associated with the background Hoescht staining in the cytoplasm.

Next, we performed unsupervised hierarchical gene clustering based on highly correlated phenotypic features, as signified by proximity in dendrograms (Supplementary figure 10; explorable in the ‘Supplementary phenotypic heatmap’ file and Supplementary table 7). This process revealed clusters that were related to the same protein complexes highlighted by the UMAP plot. The ‘dynein-dynactin’ cluster (Figure 4C) consisted of neighbouring sub-clusters of genes encoding a subset of dynein (*DYNC1H1, DYNC1I2, DYNLRB1*) and dynactin (*ACTR10, DCTN6, ACTR1A, DCTN5*) constituents.

Also present in the dynein-dynactin cluster were *FAM160A2, AKTIP* and *HOOK3* (which themselves formed a sub-cluster), *DYNLL1, LIS1*, and *BICD2*, as well as six additional genes. These six genes were *SUGP1* (also known as *SF4*), which encodes an RNA-binding protein with a remarkably similar phenotypic fingerprint to that of the dynein and dynactin components, a grouping of *FLCN, RBM48* and *TMED2* (which encode a GTPase-activating protein, RNA-binding protein and transmembrane protein, respectively), as well as the stress sensor *NRF1* and a protein generated by readthrough between the genes encoding the E3 ligase RNF103 and the multivesicular body component CHMP3 (*RNF103-CHMP3*), which were both grouped with *FAM160A2, AKTIP* and *HOOK3*.

However, not all components of dynein and dynactin identified in the screen were present in the dynein-dynactin cluster. The phenotypic signatures of *DYNC1LI1* and the dynactin component *CAPZB* were divergent from those of other known dynein and dynactin components, as well as from each other (Supplementary figures 10 and 11). This could be because these factors take part in a specific subset of motor functions or have additional, dynein-independent functions that influence cellular organisation. Consistent with this notion, the DYNC1LI1 and DYNC1LI2 light intermediate chains have non-overlapping functions in some dynein-based trafficking events ^70^ and CAPZB has an additional role in capping F-actin ^71^. Nonetheless, our analysis shows that phenotypic clustering is a valuable tool for revealing novel gene associations in our dataset, as well as for highlighting factors that are strong candidates to work closely with the dynein complex.

### Hit verification with independent crRNAs

To evaluate crRNA specificity in the aforementioned experiments, we targeted a subset of hits from the secondary screen with unrelated pools of crRNAs. These reagents were selected from the Vienna Bioactivity CRISPR (VBC) collection, which referentially targets functional protein domains ^72^. For these experiments, we selected five of the six additional genes that clustered with dynein-dynactin components, as well as 17 other genes whose targeting caused mislocalisation of at least one dynein cargo.

As in the secondary screen, we assessed effects of the independent crRNA pools on subcellular localisation of GFP-BICD2N-FRB and PTS-RFP-FKBP in U-2 OS PEX cells treated with rapamycin, as well as EEA1 and TGN46 in untreated, unmodified U-2 OS cells. We also included a new readout of dynein activity by staining the untreated, unmodified cells with an antibody to LAMP1, which marks lysosomal compartments. These structures have previously been shown to rely on dynein for trafficking towards the perinuclear region ^54, 73, 74^ and we corroborated this conclusion in U-2 OS cells using *crDYNC1H1* and *crLIS1* (Supplementary figure 7).

Significant effects on cargo localisation were confirmed for 19 out of the 22 hits (86.4%) using the VBC crRNA pools (Figure 5 and Supplementary figure 12). The lack of activity for at least two of the other three pools (which were directed against *CDH23, FAM86B2* and *RNF183*) appeared to reflect inefficient cutting of the target gene (Supplementary figure 13). Overall, however, these experiments revealed a high rate of hit replication.

**Figure 5.**
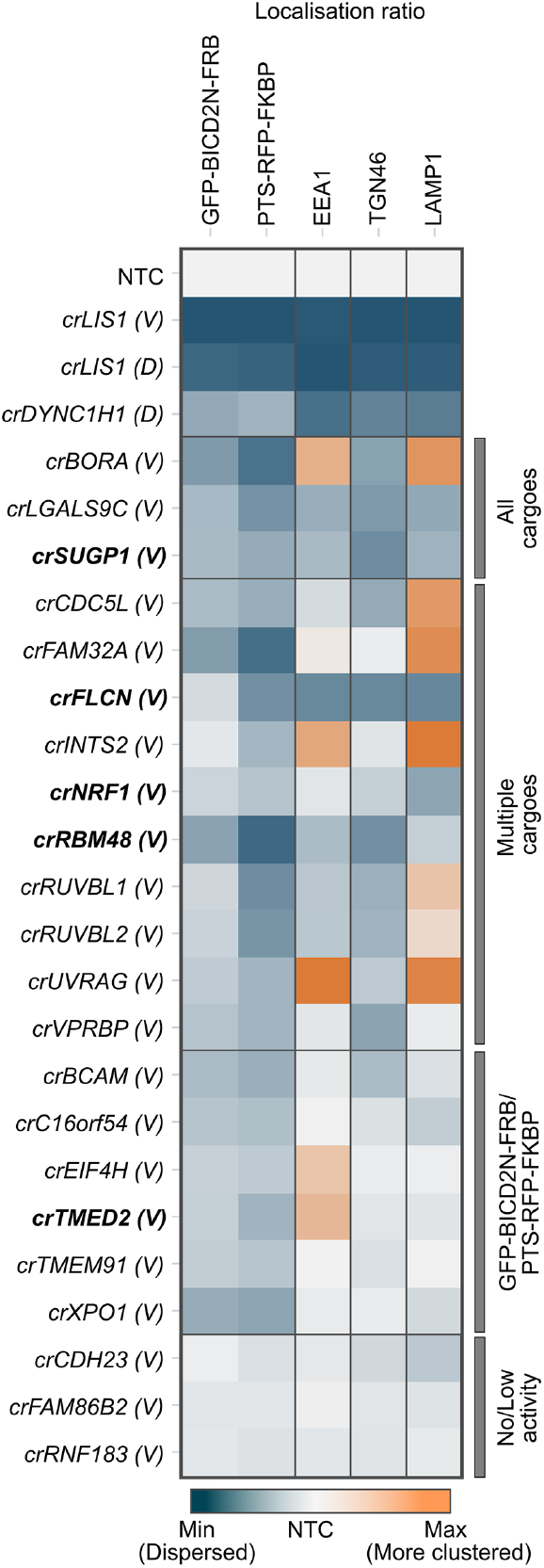
Orthogonal cargo localisation screen with a subset of candidates. Heatmap displaying localisation ratio of dynein cargoes at the perinuclear vs peripheral region of cells treated with the indicated crRNAs. GFP-BICD2N-FRB and PTS-RFP-FKBP were evaluated in U-2 OS PEX cells treated with rapamycin, whereas other markers were evaluated in unmodified, untreated U-2 OS cells. crRNAs were synthesised based on the Vienna Bioactivity CRISPR score (labelled ‘(V)’), except for *LIS1* and *DYNC1H1* crRNAs from the initial Discovery set (labelled ‘(D)’), which were used as additional positive controls. Bold labelling indicates crRNAs that target novel constituents of the ‘dynein-dynactin’ cluster previously generated by unsupervised profiling. Colour scales of individual features were adjusted based on their min and max values. Categories of affected cargoes were manually annotated based on statistically significant effects (see Supplementary figure 12). Data represent relative change of the mean aggregated at well level compared to NTC from a minimum of three independent experiments (minimum of 100 cells analysed per well, four wells analysed per condition).

This analysis additionally revealed that crRNA pools targeting *NRF1, SUGP1, FLCN* and *LGAL9SC* caused dispersion of lysosomal compartments (Figure 5 and Supplementary figure 12). Moreover, we found that, in addition to reducing the enrichment of GFP-BICD2N-FRB and PTS-RFP-FKBP to the perinuclear region, crRNAs to *CDC5L* and *FAM32A* enhanced clustering of lysosomal compartments at this site. crRNAs to *BORA, INTS2* and *UVRAG* increased perinuclear clustering of both lysosomal compartments and early endosomes, whilst causing dispersion of peroxisomes (Figure 5 and Supplementary figure 12). These observations raise the possibility of interplay between different dynein-based trafficking processes (see Discussion).

### SUGP1 sustains functional levels of LIS1 mRNA

Finally, we investigated the mode of action of *SUGP1*. Targeting this gene caused dispersion of all cargoes tested, as well as similar effects to known components of the transport machinery on several features related to the nucleus and microtubule cytoskeleton. Consequently, *SUGP1* grouped very closely with known components of the dynein-dynactin machinery in our phenotypic clustering analysis (Figure 4C). *SUGP1* encodes a 72-kDa protein that contains two SURP domains and a G-patch domain (Figure 6A).

**Figure 6.**
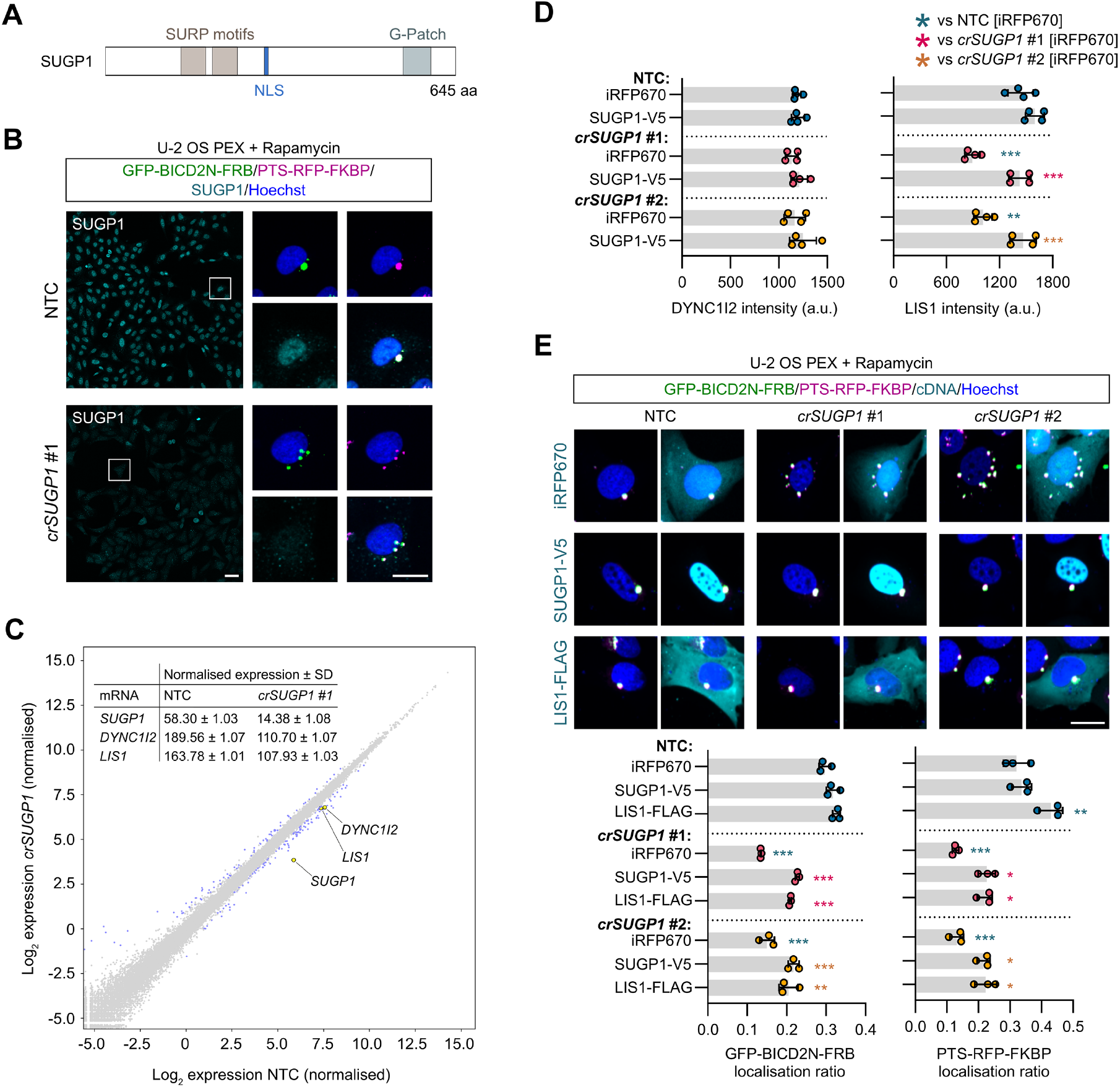
SUGP1 sustains functional levels of *LIS1* mRNA and protein. **A)** Schematic of SUGP1 domain structure. NLS, nuclear localisation signal. **B)** Representative images of SUGP1 intensity and GFP-BIC2N-FRB and PTS-RFP-FKBP localisation in U-2 OS PEX cells treated with *crSUGP1* #1. Scale bar, 25 μm. **C)** Scatter plot of mRNA abundance for *crSUGP1* #1-edited vs NTC U-2 OS cells (mean log_2_ normalised values from three independently performed experiments). mRNAs meeting the threshold for inclusion (minimum absolute log_2_ fold change ≥ 0.5; FDR ≤0.05) are labelled in blue, except *SUGP1, DYNC1I2* and *LIS1*, which are labelled in yellow. Inset table shows non-logarithmic values for *SUGP1, DYNC1I2* and *LIS1*. See Supplementary table 8 for full results. **D)** Quantification of endogenous DYNC1I2 and LIS1 protein signal (assessed by immunofluorescence) in unmodified U-2 OS treated with NTC, *crSUGP1* #1 *or* #2 and transfected with a control (iRFP670) or crRNA-resistant SUGP1-V5 expression plasmid. **E)** Representative images and quantification (perinuclear vs peripheral localisation ratio) of GFP-BICD2N-FRB and PTS-RFP-FKBP localisation in U-2 OS PEX cells treated with NTC, *crSUGP1* #1 or #2 and transfected with a control (iRFP670), crRNA-resistant SUGP1-V5, or LIS1-FLAG expression plasmid. Scale bar, 25 μm. In D and E, cells were transfected with crRNA 96 h before fixation, and with expression plasmid 48 h after crRNA transfection. Data points are aggregated means of independent experiments, with a minimum of 100 transfected cells analysed per condition. Error bars signify S.D.. \**p*<0.05, \*\**p*<0.01, \*\*\**p*<0.001 (two-way ANOVA with Tukey’s multiple comparison; colours of asterisks indicate comparison group).

These motifs are found in many eukaryotic RNA-processing enzymes, leading to annotation of SUGP1 as an RNA-binding protein. Little is known about the molecular and cellular functions of the protein, other than roles in SF3B1-associated 3′-splice site recognition during pre-mRNA processing ^75, 76^ and regulation of cholesterol metabolism ^77^.

We first corroborated the requirement for SUGP1 in dynein-based trafficking by revealing inhibitory effects of seven out of eight individual crRNAs on both SUGP1 protein expression and peroxisome relocalisation in U-2 OS PEX cells (Figure 6B and Supplementary figure 14A). As SUGP1 is enriched in the nucleus (Figure 6B) and is a predicted RNA-binding protein, we hypothesised that its depletion affects expression or processing of components of the dynein complex or its regulators. To test this notion, we performed RNA-seq analysis of polyA-enriched RNAs in U-2 OS cells edited with a single, highly active *SUGP1* crRNA (Figure 6B and Supplementary figure 14A). As controls, we processed cells treated with a crRNA from the NTC pool or a crRNA that targets the *XCR1* gene. Because *XCR1* is exclusively expressed in a subset of dendritic cells ^78^, its targeting should report on any changes in the U-2 OS transcriptome that are induced purely by DNA double-strand breaks. We confirmed that the selected *SUGP1* and *XCR1* crRNAs edited the target sites in the samples processed for RNA-seq (Supplementary figure 14B – D) and that the *XCR1* crRNA had no effect on dynein activity in the peroxisome relocalisation assay (Supplementary figure 14A).

Only a single transcript, *XIRP1*, met our criteria for a significant change in abundance in NTC vs *crXCR1* samples (minimum absolute log2 fold change ≥ 0.5 and false discovery rate ≤ 0.05; Supplementary figure 15A and Supplementary table 8). In contrast, the abundance of 149 mRNAs was significantly altered in *SUGP1*-edited cells compared to both NTC and *crXCR1* controls (Figure 6C, Supplementary figure 15B, and Supplementary table 8). This list included two mRNAs encoding known components or regulators of the dynein machinery – *DYNC1I2* and *LIS1* – which exhibited, respectively, ∼40% and ∼35% reductions in abundance when compared to the controls. By comparison, *SUGP1* mRNA levels were reduced by ∼75% compared to the controls. The changes in *DYNC1I2* and *LIS1* mRNA levels were independently validated by RT-qPCR in both U-2 OS cells and ARPE-19 cells treated with a different *SUGP1* crRNA (Supplementary figure 15C). We conclude from these experiments that levels of *DYNC1I2* and *LIS1* mRNA are reduced in *SUGP1*-edited cells.

Further analysis of the RNA-seq data with the rMATs pipeline ^79^ revealed 189 genes with significantly altered splicing patterns in *SUGP1*-edited cells compared to both controls (Supplementary figure 16A – C and Supplementary table 9), whilst the LABRAT package ^80^ showed that 17 genes had 3′-end usage that was significantly affected by *SUGP1* targeting (Supplementary figure 16D and Supplementary table 10). However, no known components or regulators of the dynein machinery exhibited altered splicing or 3′-end usage (Supplementary tables 9 and 10). Thus, the reductions in *DYNC1I2* and *LIS1* mRNA level in *SUGP1*-edited cells appear not to be associated with changes in splicing or alternative polyadenylation of these transcripts.

We next investigated the consequences of reduced *DYNC1I2* and *LIS1* mRNA levels in *SUGP1*-edited cells on the abundance of their protein products using immunofluorescence. The DYNC1I2 protein signal was not altered by *crSUGP1* (Figure 6D), revealing the existence of a mechanism that compensates for lower concentration of its mRNA. LIS1 protein signal was, however, significantly reduced when SUGP1 was disrupted with crRNA (29.5 and 38% reduction for two different SUGP1 guides compared to NTC) (Figure 6D). This deficit in LIS1 levels could be fully restored by transfection of a crRNA-resistant *SUGP1* cDNA (Figure 6D). These observations indicate that SUGP1 promotes the expression of LIS1 protein by controlling the level of its mRNA.

Given that even a small reduction in LIS1 protein abundance can have significant functional consequences ^17, 20, 21, 23, 81, 82^, our findings led us to hypothesise that lowered LIS1 levels contribute to dynein trafficking defects in *SUGP1*-edited cells. If this notion were correct, increasing LIS1 levels should suppress the cargo mislocalisation phenotype. To test if this is the case, we transfected *SUGP1*-edited and NTC U-2 OS PEX cells with a *LIS1* cDNA and treated them with rapamycin. In NTC cells, the *LIS1* cDNA elicited no change in GFP-BICD2N-FRB distribution and only a very modest increase in clustering of the PTS-RFP-FKBP signal compared to controls in which a cDNA encoding the fluorescent protein iRFP670 was transfected (Figure 6E). In contrast, *LIS1* cDNA transfection boosted perinuclear localisation of both GFP-BICD2N-FRB and PTS-RFP-FKBP in *SUGP1*-edited cells (Figure 6E). The magnitude of the suppression of the mislocalisation phenotype was very similar to that observed in *crSUGP1* cells transfected with the *SUGP1* cDNA (Figure 6E). These data indicate that a key function of SUGP1 in the context of dynein-based trafficking is sustaining *LIS1* mRNA levels.

## DISCUSSION

Our knowledge of dynein regulation is currently limited to studies of a small number of binding partners, most notably dynactin, LIS1 and a handful of activating adaptors. To gain new insights into dynein biology, we have performed a genome-wide loss-of-function CRISPR screen in human cells using the subcellular distribution of the motor’s cargoes as a functional readout. Our findings represent a valuable resource for phenotypic analysis of dynein-based trafficking, as well as other aspects of cellular organisation.

### Leveraging arrayed CRISPR screening to reveal requirements for dynein-based trafficking

Genome-wide CRISPR screening drives new biological discoveries by unbiased interrogation of gene function. The vast majority of CRISPR screens in the literature are in a pooled format, in which a crRNA library is introduced into a cell population en masse, followed by selection for a phenotype and identification of associated crRNAs by next generation sequencing ^83^. Whilst this method has the advantage that crRNA delivery is straightforward, it is typically limited to readouts based on chemical selection or flow cytometry and therefore is not well suited for assessing subcellular phenotypes. To address this issue, innovative pooled screening methods have recently been developed, involving photoconversion of cells with desirable phenotypes followed by cell sorting ^43, 84^, direct sorting of cells based on spatial phenotypes ^85^, and *in situ* sequencing of crRNAs ^45, 86, 87^. Nevertheless, the arrayed format remains the most direct approach for establishing genotype-phenotype associations, and is well-suited for complex imaging-based endpoints ^30, 31, 44, 88-90^. However, this method is costly in terms of upfront investment in arrayed crRNA libraries, automation, and assay development.

Here we sought to maximise the output of the arrayed screening format by optimising gene disruption procedures, including through the development of a robust and scalable protocol for mRNA-based delivery of Cas9, and multivariate analysis of the distribution of two dynein cargoes that depend on distinct activating adaptors – BICD2N-tethered peroxisomes and HOOK-associated early endosomes. The effectiveness of the screening and analysis pipeline was demonstrated by recovery of core components of the dynein transport machinery, as well as known adaptors between the motor complex and early endosomes. However, 12 of 23 (52%) components that comprise the core dynein-dynactin machinery were not identified as hits for either cargo. This observation indicates that not all genes that contribute to dynein function were identified in the screen. This is presumably due to inherent limitations of high-throughput genetic screens – functional redundancy between paralogues, perdurance of long-lived proteins following gene disruption, or some genes not being targeted efficiently or not being expressed in the studied cell type. Nonetheless, we validated 195 novel hits from the primary screen, which are candidates for future mechanistic analysis.

### Phenotypic clustering reveals novel gene associations

Using morphological profiling, we generated fine-grained phenotypic signatures of the hits from the secondary screen. This strategy has previously been used to assess cell state ^91^, predict the underlying biological activity of small molecules ^92^, and identify co-functional genes in functional genomics data ^45, 93-95^. We found that features associated with just seven markers were sufficient to cluster components of protein complexes with diverse functions, including dynein-dynactin. We also observed clustering of six genes with dynein-dynactin that do not encode components of the core machinery. Of these genes, four had not previously been linked physically or functionally with the motor complex. The importance of three of these factors – *RBM48, SUGP1* and *TMED2* – for cargo distribution was confirmed with independent crRNAs. The other two genes that cluster with dynein and dynactin components were *NRF1* and *FLCN. NRF1*, targeting of which caused mild peroxisome and lysosome dispersion phenotypes, encodes a transcriptional factor that can complex with DYNLL1 and DYNLL2 ^96^. However, it is not known if these interactions are related to the light chains’ function in the motor complex, or their dynein-independent role in assembling protein complexes ^97^. FLCN is a GTPase activating protein that promotes retrograde trafficking of lysosomes by dynein ^74, 98-100^, potentially by stabilising the interaction of the activating adaptor RILP with Rab34 ^101^. We found that not only lysosomes, but also early endosomes and the Golgi, were dispersed when *FLCN* was targeted. As RILP function appears to be restricted to late endosomes, lysosomes and autophagosomes ^74, 99, 102, 103^, our findings raise the possibility that FLCN has a RILP-independent function in trafficking of other dynein cargoes. However, FLCN does not appear to have a general function in dynein-based trafficking as its disruption did not affect BICD2N-mediated localisation of peroxisomes.

Clearly, the hits from the dynein-dynactin cluster should be prioritised for mechanistic studies. However, whilst many of the genes that do not fall in the dynein-dynactin cluster will indirectly influence the localisation of cargoes, we anticipate that a subset will directly influence motor function. This view is supported by our observation that the phenotypic profiles of the Dynein light intermediate chain isoform *DYNC1LI1* and the dynactin component *CAPZB* do not cluster with those of other dynein subunits. As discussed earlier, factors with direct effects on the dynein machinery could have divergent phenotypic signatures because they affect a small subset of motor functions and/or have additional, dynein-independent functions that also alter cellular organisation. Studying hits from outside the dynein-dynactin cluster could, therefore, also shed light on transport mechanisms. A strong candidate to follow up in this regard is *LGALS9C*. crRNAs targeting this protein, which codes for a galectin-9 isoform, disrupt localisation of all dynein cargoes tested and have a phenotypic profile that clusters with that of *CAPZB*.

### Selectivity, interplay and competition during cargo trafficking

We observed a number of genes whose targeting altered localisation of only a subset of dynein cargoes. Mechanisms that could account for such a phenomenon include modulating the expression, localisation or function of a subset of cargo adaptors or regulating the activity of dynein only when it is engaged with certain cargoes. One hit in this category was the key nuclear export protein XPO1 (also known as Exportin-1 and CRM1). Disruption of XPO1 is known to affect several microtubule-related events, including spatial organisation of MTOCs ^104^, recruitment of proteins to the centrosome ^105^ and perinuclear detachment of dynein-associated adenovirus from microtubules ^106, 107^. In our experiments, targeting XPO1 caused dispersion of peroxisomes but not other dynein cargoes tested. These data suggest that this protein is not required for all microtubule-based processes.

Unexpectedly, we also found that disrupting some genes increases perinuclear clustering of at least one type of dynein cargo, whilst dispersing at least one other. These factors might influence the recruitment of limiting components of the transport machinery to one cargo type, thereby affecting their availability for others. One gene in this category is *INTS2*, which codes for a nuclear protein that is best known as a mediator of small nuclear RNA processing ^108^ but also promotes recruitment of dynein to the cytoplasmic face of the nuclear envelope ^109^. We observed increased perinuclear clustering of endosomes and lysosomes when *INTS2* was targeted, which could conceivably arise from dyneins that otherwise would be located at the nuclear surface becoming accessible for other cargoes. Collectively, these observations raise the possibility that activities of different dynein-driven processes are finely balanced. Competition for components of the transport machinery is also likely to explain the finding that disrupting the acetylhydrolase PAFAH1B2 increases clustering of endosomes near the nucleus. This finding adds to evidence that PAFAH1B2 sequesters LIS1 from dynein and thereby influences motor activity ^48, 110^. Whether there are active mechanisms that regulate shuttling of LIS1 between PAFAH1B2 and dynein warrants further investigation.

### SUGP1 has a general role in dynein-based trafficking and sustains LIS1 levels

To demonstrate the utility of our screen for generating novel mechanistic insights, we investigated the mode of action of the RNA-binding protein SUGP1. This factor had a phenotypic signature that correlates closely with components of the dynein complex. We found that SUGP1 disruption lowers *LIS1* mRNA and protein levels, and that restoring LIS1 protein levels suppresses the associated defects in cargo localisation. Whilst we cannot rule out additional roles of SUGP1 in dynein-based trafficking, our experiments suggest a key function of this protein is promoting expression of LIS1. Although the reduction of LIS1 protein in SUGP1-edited cells was modest, lowering LIS1 abundance by a similar amount is sufficient to impair dynein-mediated trafficking and cause neurodevelopmental defects in animal models ^17, 22^. In keeping with these findings, loss of one copy of the *LIS1* gene causes severe brain malformation in humans ^21, 23, 81, 82^. As the splicing and 3′-end usage of *LIS1* mRNA was not altered by disruption of SUGP1, the protein presumably plays a role in transcription or stabilisation of *LIS1* mRNA, or affects processing of mRNAs whose products control these events. Further investigation of how SUGP1 promotes *LIS1* expression could shed light on post-transcriptional regulation of dynein-based trafficking, which is a fascinating but largely unexplored topic.

### A resource for shedding light on other aspects of cellular organisation

Whilst our main objective has been to reveal genetic requirements for dynein-based trafficking, we anticipate that the output of this project will be valuable for those interested in several other aspects of cell biology. Our primary screen additionally highlighted crRNAs that affect the morphology of early endosomal compartments and peroxisomes without altering their positioning. We also identified novel factors that influence nuclear morphology, formation or micronuclei, organisation of the microtubule network, or the number of MTOCs. Our unbiased phenotypic clustering of hits from the primary screen additionally revealed many novel gene associations, including those involving known players in the regulation of gene expression and protein homeostasis. Exploring these associations is likely to provide new insights into the molecular control of these processes. Finally, the images from the primary screen can be mined to extract additional spatial phenotypes and identify genes that affect them.

## MATERIALS AND METHODS

### Cell culture

The U-2 OS human bone osteosarcoma cell line stably expressing GFP-BICD2N-FRB and peroxisome targeting sequence (PTS)-RFP-FKBP was previously described ^27^. U-2 OS, HEK-293, and IMR-90 cells were maintained in McCoys 5A, DMEM and Eagle’s MEM, respectively, whereas ARPE-19 and SH-SY5Y cells were cultured in DMEM/Nutrient Mixture F12 Ham. All media was purchased from Sigma Aldrich and was supplemented with 10% (v/v) FBS and 1% (v/v) GlutaMAX (Thermo Fisher). All cell lines were certified free of Mycoplasma either internally using the MycoSEQ Mycoplasma detection kit (Thermo Fisher) or by IDEXX BioAnalytics using STAT-Myco testing. The identities of cell lines were authenticated by short tandem repeat fingerprinting by IDEXX BioAnalytics. 3

### mRNA-Cas9 transfection

For optimisation of mRNA delivery, cells were seeded at ∼70% confluency and reverse transfected with indicated concentrations of mRNA-Cas9-HA containing 5-methoxyuridine (5moU) or mRNA-eGFP 5moU (TriLink) in MessengerMAX (Thermo Fisher) or RNAiMAX (Thermo Fisher). Samples were fixed either 6- or 24-h post-transfection and processed for immunofluorescence. Cas9 expression was monitored by the intensity of HA signal, which was quantified using Columbus 2.9.1 (Perkin Elmer). For screening and all other experiments, the concentration of mRNA-Cas9-HA was fixed at 40 ng per well of a 384-well plate. Cells were reverse transfected with mRNA-Cas9-HA coupled with 1% (v/v) MessengerMAX in OptiMEM for 6 h prior to crRNA transfection.

### Synthetic crRNA preparation, transfection and automated liquid handling

Synthetic two-part crRNA (synthesised by Horizon Discovery) was used for all experiments. The whole-genome human crRNA library was prepared as described previously ^31^, with pools of four equimolar crRNAs targeting each gene. Controls included in the genome screen were: NTC (non-targeting controls; a pool of four pre-designed crRNAs (Horizon Discovery) with at least three mismatches to potential PAM-adjacent targets in the human genome; 38 wells per plate) as the neutral control; *crLIS1* (a pool of four crRNAs targeting *LIS1*; 13 wells per plate) as the positive control, and *crPLK1* (a pool of four crRNAs targeting *PLK1*) as the editing control. For hit confirmation, identical pools of crRNAs (n = 384 genes) and/or a new pool of four crRNAs per gene selected based on the Vienna Bioactivity CRISPR score ^72^ (VBC; n = 22 genes) were custom synthesised by Horizon Discovery. In the secondary screens, individual NTC crRNAs (NTC 1 – 4), the *crLIS1* pool and *crPLK1* pool were included in the synthesis of the library plate as additional controls. These were used for quality control of individually synthesised batches of crRNAs via assessment of their performance in the phenotypic assays. crRNAs were dispensed into 384-well Phenoplates (Perkin Elmer) via an Echo 555 instrument (Labcyte) at a final concentration of 50 nM. For the genome-wide screen, automatic liquid handling was executed by docking a Steristore (HighRes), Echo 555, Multidrop combi (Thermo Fisher) and washer dispenser EL406 (Biotek) to a Star6 automation platform (HighRes). Library and assay plates were stored in the Steristore at 8°C during dispensing and were brought to room temperature before reverse transfection. Cas9-transfected U-2 OS cells (1500 cells per well of the 384-well plate) were reverse transfected with crRNAs with 1% (v/v) RNAiMAX (Thermo Fisher) in serum-free media dispensed via a Multidrop combi. Plates were incubated for 72 h before rapamycin treatment, fixation and immunostaining (all via automatic liquid handling).

### Drug treatment

To induce heterodimerisation of GFP-BICD2N-FRB and PTS-RFP-FKBP, cells were (unless indicated otherwise) treated with 2 nM rapamycin (SelleckChem) in serum-free media via a Multidrop combi for 2.5 h before fixation. Nocodazole (3 μM; Sigma Aldrich) and DMSO as vehicle control were added via a Tecan D3000 dispenser.

### Immunostaining

Cells were fixed in 4% (w/v) paraformaldehyde containing 0.04 mg/mL (w/v) Hoechst 33342 (Thermo Fisher) for 20 min, followed by permeabilisation and blocking in 5% (w/v) bovine serum albumin (BSA) and 0.25% (v/v) Triton-X-100 in phosphate-buffered saline (PBS) for 1 h. Cells were then were incubated in 1% (w/v) BSA with primary antibodies at 4°C overnight followed by secondary antibodies at room temperature for 2 h (see Suppementary table 11 and 12 for details of primary and secondary antibodies, respectively). All washes were performed with PBS on a Washer Dispenser EL406 (Biotek). For the secondary screen, U-2 OS PEX cells were stained with α-Tubulin antibodies and Hoescht (in addition to directly visualising GFP-BICD2N-FRB and PTS-RFP-FKBP), whereas unmodified U-2 OS cells were stained with TGN46, EEA1 and γ-Tubulin antibodies, as well as Hoescht.

### High-content imaging, feature calculations and data normalisation

Image acquisition was performed on the CellVoyager 8000 (Yokogawa) with either a 20x or 40x water immersion objective (1.0 and 0.95 NA, respectively) with a minimum of four fields of view per well. Images were acquired as complete Z-stacks and were post-processed to generate maximum projections. 2×2 binning was performed on the images acquired for the genome screen to reduce file size. All analysis was performed on binned 20x images except for the analysis on α-tubulin in the secondary screens, which was performed on unbinned 40x images. Cellular segmentation and image analysis were performed using Columbus 2.9.1 (Perkin Elmer). In brief, nuclei were detected by virtue of strong Hoechst signal, whereas cytoplasm was detected using either the diffuse, background Hoechst signal or the outline of the α-tubulin antibody signal. Nuclei were subsequently filtered based on cellular morphology, with those located at the image boundary excluded.

For the peroxisome relocalisation assay, nuclei were further selected for GFP- and RFP-positive cells based on their intensity profiles. Calculations of morphological features or spot quantification were performed with pre-existing Columbus algorithms. To determine the ‘localisation ratio’ of cargoes, the detected cells were segmented into two rings – the perinuclear region (with an outer limit of 7 μm from the nuclear envelope) and peripheral region (> 7 μm from the nuclear envelope) – for spot detection. A cut-off of 7 μm was selected based on pilot studies that compared cargo distribution in control U-2 OS cells with those treated with nocodazole. Localisation ratio was determined by dividing the number of spots in the perinuclear region by the number in the peripheral region. Where specified, the peripheral region was further segmented into intermediate (with a distance of 7 μm to 14 μm from the nuclear envelope) and outer regions (with a distance of > 14 μm from the nuclear envelope). A minimum of 100 cells per well were analysed, with mean values per well used for downstream analysis.

Data normalisation and transformation including linear discriminant analysis (LDA) was performed with Genedata Screener 19.0.1 (Genedata). LDA was used as a linear classifier by combining multiple individual features (that have an rZ ≥ 0.1) to facilitate separation of the NTC versus a positive control and/or to generate a single feature for hit calling. LDA was also used for morphological and/or textural analysis when no individual feature was sufficient to define the phenotype of interest. Either one-point normalisation (rZ-score with NTC as a central reference) or two-point normalisation (NTC minus positive control; 0 – 100) was used. For the genome-wide screen, genes were initially filtered based on number of cells, cellular morphology and micronuclei before hit calling. Thresholds were based on mean ± S.D. and were assessed and adjusted based on the desired stringency (at least ± 2*S.D. or more) compared to the specific control (*e*.*g*. NTC) for individual features. The rationale for using a range of thresholds rather than a fixed threshold in the genome-wide screen was to take into consideration the limitations of the one-shot screening (*i*.*e*. with no replicates for almost all crRNA pools), as well as the number of hits that we had the capacity to take forward to the secondary screen. For the subsequent hit validation activities, the increased number of technical and biological replicates provided more confidence when hit calling and therefore we standardised on a minimum threshold of at least ± 2.5* S.D. of NTC. Thresholds used in specific instances are given in the figure legends.

### Image-based profiling

Mean profiles of features (n = 2003) were generated with the pre-existing algorithms in Columbus 2.9.1 (Perkin Elmer). Data were transformed into rZ-scores with NTC as a central reference, and were aggregated by median per crRNA pool. Feature reduction was performed by removing highly variable features (defined by S.D. ≥ 3 of *crLIS1* controls that were scattered around the 384-well plates) and highly-correlated (*i*.*e*. redundant) features (*R*^2^ ≥ 0.8). The remaining features (n = 278) were subjected to hierarchical clustering to group crRNAs using complete linkage and correlation for distance measures with a normalisation of scaling between 0 and 1.

### Molecular cloning and DNA transfection

The cDNA sequence for human SUGP1 (based on RefSeq: NM_172231) fused with a C-terminal V5 epitope tag was synthesised and cloned into the KpnI and XbaI restriction sites in pcDNA3.1(+) by Azenta Biosciences. The synthesised *SUGP1* sequences had mutated PAM or target site sequences to prevent cutting of the cDNA by *SUGP1* crRNAs in rescue experiments. The expression construct for human LIS1 (RefSeq: NM_000430) fused with a C-terminal FLAG epitope tag was obtained from OriGene. An iRFP670 expression plasmid (synthesised by Azenta Biosciences) in the pcDNA3.1 (+) backbone was used as a transfection control. Cells were transfected with endotoxin-free plasmid DNA 48 h post-crRNA transfection and incubated for an additional 48 h before fixation. DNA transfection was performed with Lipofectamine 3000 (Thermo Fisher) according to the manufacturer’s instructions (for each well of a 384-well plate: 6 ng plasmid DNA, 1% [v/v] Lipofectamine 3000, 0.5% [v/v] p3000 reagent). Transfected cells were selected based on the intensity profile of either iRFP670, V5 or FLAG.

### Tracking of Indels by DEcomposition (TIDE)

Genomic DNA was isolated from cells collected from 96-well plates using Direct PCR lysis reagent (Viagen) supplemented with 1 mg/mL (w/v) Proteinase K (Sigma Aldrich) according to the manufacturer’s instructions. PCR reactions were carried out with Phusion Flash PCR Master Mix (Thermo Fisher), again as instructed by the manufacturer. PCR products were purified using the QIAquick kit (Qiagen) and capillary sequencing was performed by Azenta Biosciences. Sequencing traces from cells treated with crRNAs targeting the gene of interest or from NTC controls were analysed using the standard parameters of the TIDE webtool ^111^.

### RNA sequencing and bioinformatic analysis

Approximately 1×10^6^ U-2 OS cells per sample were harvested from an individual T25 flask and processed for RNA extraction and sequencing. The samples were from three independent experiments that each included U-2 OS cells transfected with pre-designed commercially available NTC guide 1, *crSUGP1* guide 1 or *crXCR1* guide 1 (Horizon Discovery). To confirm editing by the guides, DNA from a proportion of cells from each sample pool was purified and analysed using TIDE, as described above. To confirm depletion of SUGP1 protein specifically in *crSUGP1*-treated cells, a proportion of each sample pool was re-seeded at high density in a 384-well plate and incubated for 24 h before fixation and processing for immunostaining with α-SUGP1 antibodies. Quality control on the extracted RNA was performed with a Qubit 4.0 Fluorometer (Thermo Fisher) and Agilent 3500 Fragment Analyzer (Agilent). Subsequent steps of RNA-seq were outsourced to Azenta who generated sequencing libraries from polyA-enriched RNA (captured with oligo-dT beads) using the NEBNext Ultra II RNA library prep kit for Illumina (NEB), multiplexed them, and loaded them on an Illumina NovaSeq 6000 machine. Samples were sequenced using a 2×150 pair-end configuration v15, followed by removal of adaptor sequences and poor quality sequence with the Illumina bcl2fastq program (version 2.20).

We validated the overall quality of the sequencing data using FastQC (version 0.11.5; https://www.bioinformatics.babraham.ac.uk/projects/fastqc/) and determined the absence of potential sources of contamination using the FastQ Screen tool (version 0.14.1) ^112^. We then performed further quality trimming of the sequences using TrimGalore! (version 0.6.7) (https://www.bioinformatics.babraham.ac.uk/projects/fastqc/) and its dependency Cutadapt (version 2.4) ^113^ in paired-end mode.

The FASTQ reads were mapped to the human reference genome GRCh38 (version 102) using the STAR aligner (version 2.7.9a) ^114^ that was distributed with the transcript splicing analysis software rMATS ^79^ inside a Docker container (made available on Docker Hub by the Xing group (https://hub.docker.com/r/xinglab/rmats)). The primary alignments of the mapped data (in the form of BAM files) were imported into the genome browser SeqMonk v1.48.0 (https://www.bioinformatics.babraham.ac.uk/projects/seqmonk/). Following data import, genes exhibiting an appreciable level of expression were identified using the *RNA-seq Quantitation Pipeline* in SeqMonk. The pipeline counted the number of read pairs mapping to exons for every gene. These gene-level expression values were normalised by dividing by the total number of mapped read pairs per sample. The results were then log2 transformed. Genes with an expression score of greater than -2 in at least one of the nine samples were selected for differential expression analysis. The number of read pairs mapping to each exon was determined again using SeqMonk’s *RNA-seq Quantitation Pipeline* but on this occasion the raw counts were recorded and no subsequent normalisation and log2 transformation steps were performed. The raw counts were used to identify differentially expressed genes using DESeq2 (version 1.32.0) ^115^, which was launched from SeqMonk using default settings. Only the count data from genes previously shown to exhibit an appreciable level of expression in one of the datasets were passed to DEseq2 for analysis. DESeq2 was run using R (version 4.1.0 (Camp Pontanezen) Patched (2021-06-08 r80465)). Differentially expressed genes were defined as having an adjusted p-value ≤ 0.05 after multiple testing correction and an absolute log2 fold change ≥ 0.5.

We converted the rMATs Docker container (v4.1.2) into a Singularity container (using Singularity version 3.8.0) and from within that container executed the rMATS Python script *rmats*.*py* to perform the STAR mapping and subsequent splicing analysis. The samples were mapped in paired-end mode using the default *rmats*.*py* settings. We also specified the *rmats*.*py* option *readLength* to optimise the software for sequence reads of 150 bp in length. In addition, the flag *variable-read-length* allowed the processing of reads of varying lengths, which was necessary owing to the quality trimming prior to rMATs analysis. This process produced pairwise splicing comparisons between the different conditions (NTC vs *crSUGP1, crXCR1* vs *crSUGP1* and NTC vs *crXCR1*).

Alternative polyadenylation for samples processed for RNAseq (NTC, *crXCR1* and *crSUGP1*) was quantified using LABRAT v0.3.0 (Ref. 80). LABRAT performs this analysis by quantifying the relative abundances of the last two exons of all transcript isoforms using the transcript quantification tool Salmon ^*116*^. Genes whose total abundance across all isoforms was less than 5 transcripts per million (TPM) in any sample were excluded from the analysis. False discovery rate (FDR) values were calculated using LABRAT’s linear mixed effects model followed by multiple hypothesis correction. Isoform structures and transcript sequences were derived from GENCODE v28. Genes with alternative polyadenylation were defined as having ΔΨ ≥ 0.05 (Ref. 80) and FDR ≤0.05.

### Taqman real-time PCR assay

RNA was extracted using the RNeasy mini kit (Qiagen) according to the manufacturer’s instructions. QuantiTect reverse transcriptase (Qiagen) was used for cDNA synthesis with an input of 800 ng RNA per sample, with the product diluted 1:4 in dH2O prior to Taqman assays. The reaction mix include 1X Taqman real-time PCR master mix, 1X Taqman probe(s) and cDNA template. Pre-designed Taqman probes (Thermo Fisher) were used for *LIS1* (Hs00181182_m1; FAM-MGB), *DYNC1I2* (Hs00909737_g1; FAM-MGB) and *B2M* (Hs00187842_m1; VIC-MGB) as a housekeeping control. Real-time PCR was performed in MicroAmp optical 384-well plates using the QuantStudio 6 Flex RT-PCR system (Thermo Fisher). The default thermocycling program was used (95ºC for 10 min, followed by 40 cycles of 95ºC for 15 s and 60ºC for 1 min). Data were analysed using the Quantstudio real-time PCR system software. Relative quantification (RQ) values were obtained by normalising against the expression level of NTC.

### Statistics

Statistics analyses were performed with Prism 9.3.1 (Graphpad), with types of data aggregation and tests specified in the legend.

## Supporting information

Supplementary phenotypic heatmap

## ACKNOWLEDGEMENTS

We are grateful to members of the Bullock Group at LMB, as well as the Functional Genomics and Open Innovation groups at AstraZeneca for technical support and sharing resources; in particular, we would like to acknowledge help from Morag Rose Hunter, Ceri Wiggins, Samantha Peel, James Pilling, Martin Booth, and Paul Twyman at AstraZeneca. We also thank Llywelyn Griffith (Francis Crick Institute) for advice on splicing analysis and Mariann Bienz (LMB) for feedback on the manuscript. This work was supported by the LMB-AstraZeneca BlueSky Fund (BSF31), the UK Medical Research Council (file reference numbers MC_U105178790 (to S.L.B.) and MC_UP_1201/9 (to Madeline Lancaster, who supports S.W.W.), and the National Institutes of Health (R35-GM133385, to J.M.T.). For the purpose of open access, the MRC Laboratory of Molecular Biology has applied a CC-BY public copyright licence to any Author Accepted Manuscript version arising.

## AUTHOR CONTRIBUTIONS

S.L.B. and D.R.-T. conceived the project. C.H.W., D.R.-T. and S.L.B. designed experiments. C.H.W. performed experiments. C.H.W., S.W.W., C.Q. and J.M.T. analysed data. C.H.W. and S.L.B. prepared the manuscript, which was edited and approved by all other authors.

