## Supplementary figure for "Genome-scale requirements for dynein-based trafficking revealed by a high-content arrayed CRISPR screen"

### SUPPLEMENTARY FIGURES

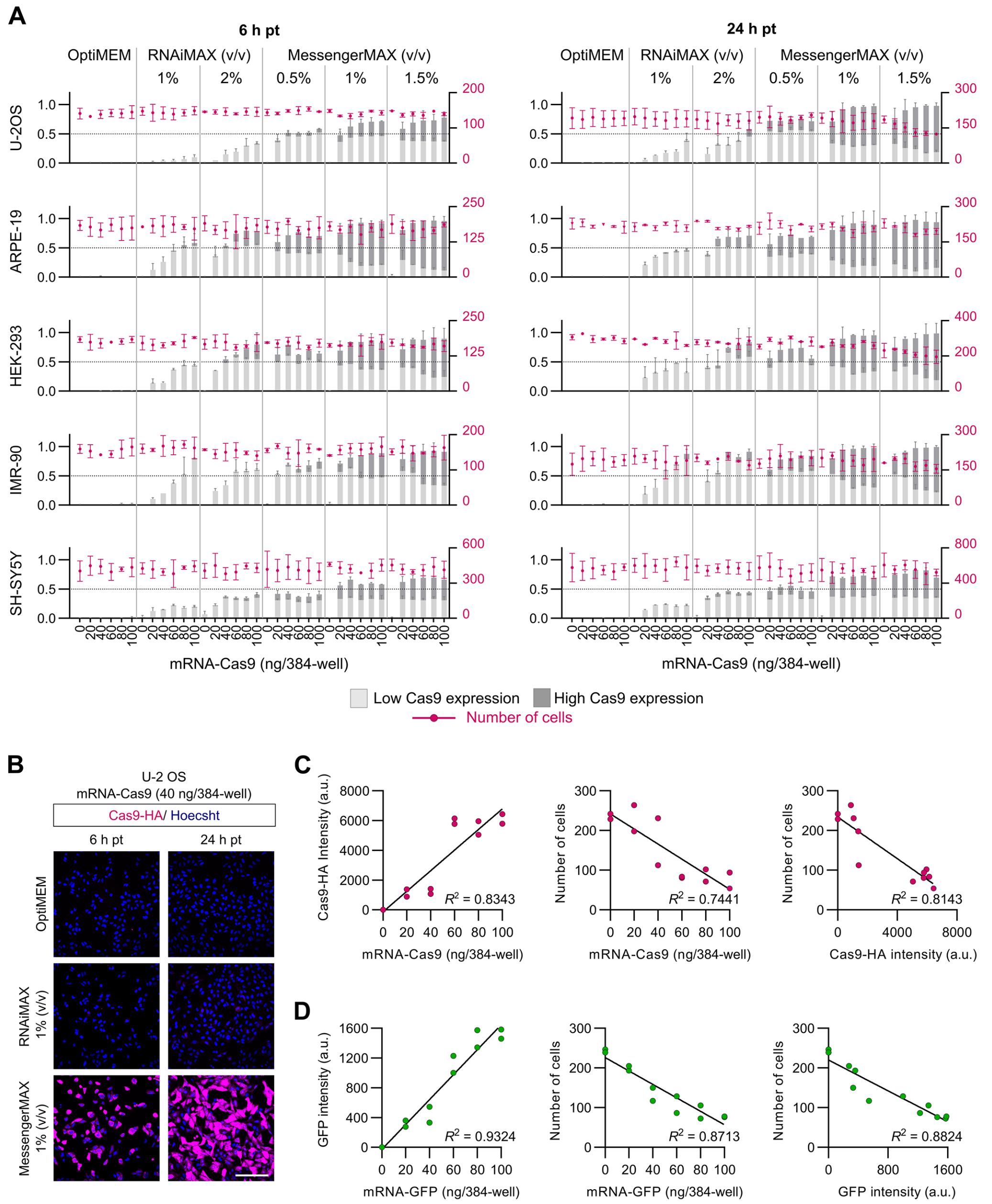


**Supplementary figure 1. Optimising conditions for mRNA-Cas9 delivery. A)** Efficiency and toxicity profile of mRNA-Cas9 delivery in a panel of five mammalian cell lines. Cells were transfected in a 384-well format with a titration of mRNA and transfection reagent and fixed either 6 or 24 h later (pt, post-transfection). OptiMEM (vehicle) was used as a control. Charts show frequency of Cas9-positive cells (immunostaining for HA tag on Cas9 with gating for low and high expression based on fluorescence intensity; bar graph, left axis, labelled in light and dark grey), and total number of cells (dot plot, right axis, labelled in magenta) as an indication of cytotoxicity. Data points represent mean aggregation at well level from two independent experiments (minimum of 100 cells from two wells analysed per condition). Error bars, S.D.. The condition selected for the study was 1% (v/v) MessengerMAX with 40 ng mRNA per well of a 384-well plate. **B)** Representative images of U-2 OS cells stained for Cas9 (HA) and DNA (Hoechst) after transfection of mRNA-Cas9 (40 ng per well of a 384-well plate) coupled with 1% (v/v) RNAiMAX or MessengerMAX. Scale bar, 200 µm. **C, D)** Reduced cell number with the highest tested MessengerMAX concentration is caused by high mRNA transfection efficiency and is not specific to Cas9 expression. U-2 OS cells were transfected with a titration of either mRNA-Cas9 or mRNA-GFP coupled with 1.5 % (v/v) MessengerMAX and fixed 24 h post-transfection. Linear regression analysis showed a negative relationship between both Cas9-HA intensity (C) and GFP intensity (D) and cell number. Data point represents mean aggregation at well level for two independent experiments (minimum of 100 cells from two wells analysed per condition).

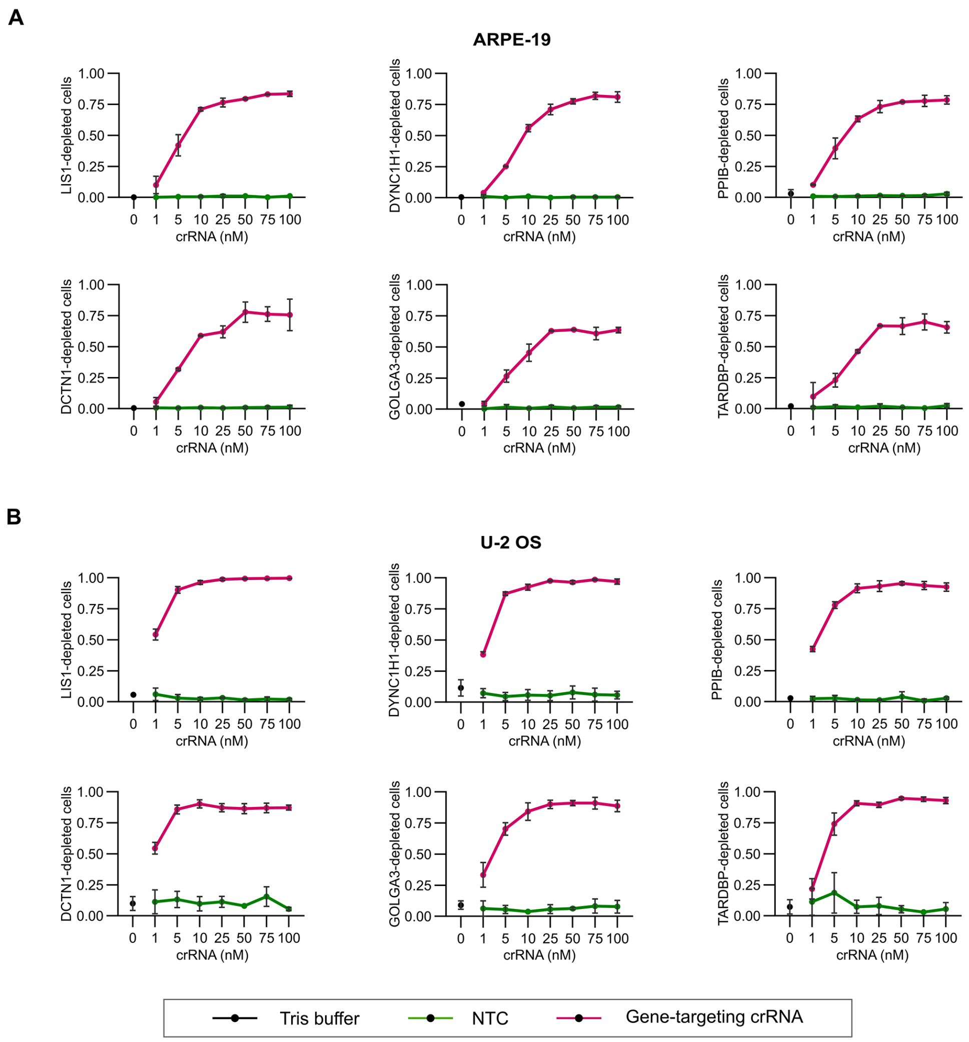


**Supplementary figure 2. Optimising crRNA concentration for CRISPR/Cas9 editing. A, B)** Quantification of CRISPR/Cas9 editing frequency in ARPE-19 (A) and U-2 OS (B) cells with varying crRNA concentration. Cells were transfected in a 384-well format with mRNA-Cas9 (40 ng/well) coupled with MessengerMAX (1%; v/v) for 6 h before transfecting with a panel of six gene-targeting crRNA pools (*crLIS1, crDYNC1H1, crPPIB, crDCTN1, crGOLGA3, crTARDBP*) or the NTC pool at the indicated concentrations. The Tris buffer used for RNA resuspension was present as an additional control. Cells were fixed 72 h later and stained with antibodies to the corresponding protein products to evaluate frequency of target protein depletion (gating performed based on the range of the protein fluorescence signal of NTC cells). Data points represent mean aggregation at well level (minimum of 100 cells from at least two wells analysed per condition). Error bars, S.D.. A final concentration of 50 nM crRNA per well was selected for the genome-wide screen.

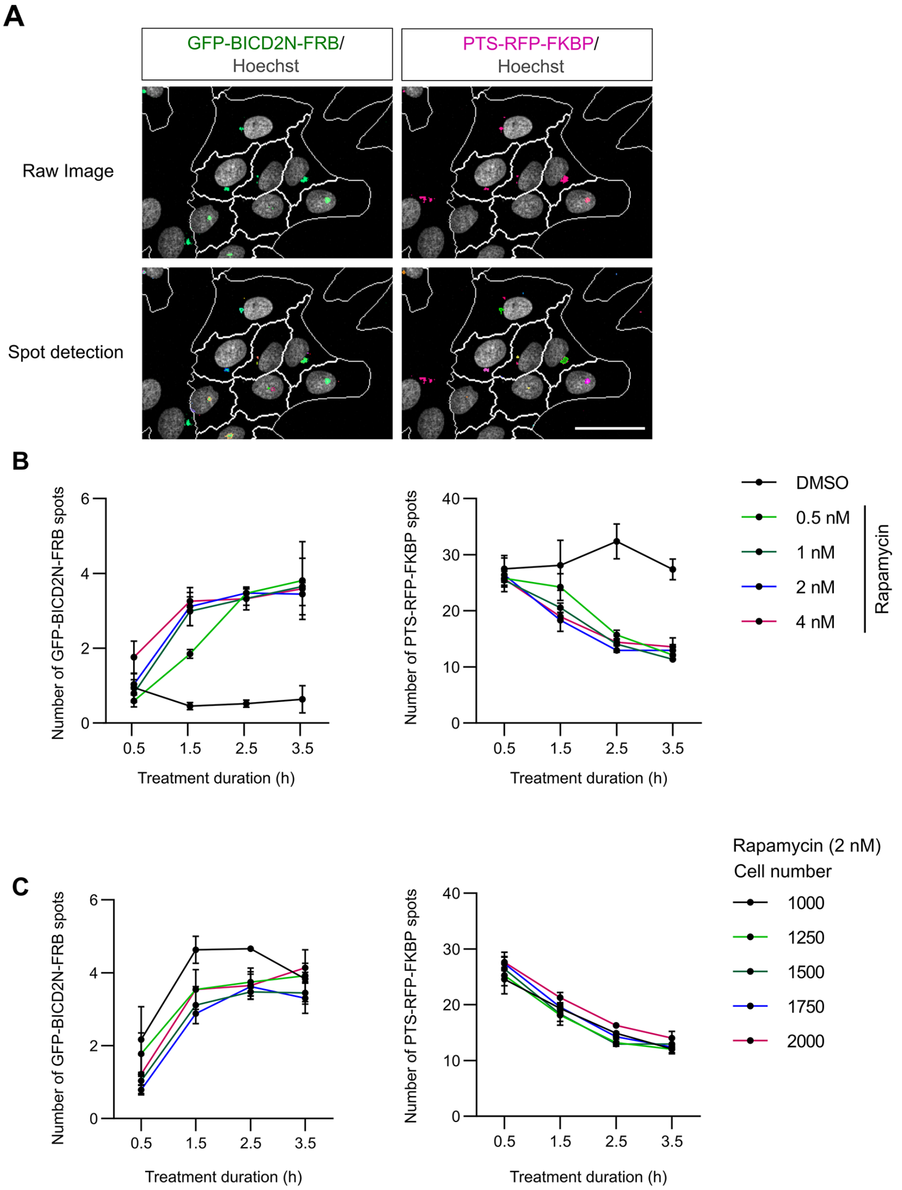


**Supplementary figure 3. Assay development for rapamycin-induced relocalisation of peroxisomes. A)** Representative images of results of applying a spot detection mask on raw images of rapamycin-treated U-2 OS PEX cells. Scale bar, 50 µm. **B)** Optimisation of rapamycin concentration and treatment duration in the peroxisome relocalisation assay. Note that, with these raw values, the number of GFP spots increases with perinuclear clustering because discrete puncta are otherwise relatively uncommon due to the dim signal, whereas RFP spot number decreases with perinuclear clustering as signals from multiple dispersed puncta coalesce at the MTOC. Data points represent mean aggregation at well level (minimum of 100 cells from at least three wells analysed per condition). Error bars, S.D.. A 2.5-h treatment with 2 nM rapamycin was selected for the genome-wide screen. **C)** Impact of number of seeded cells per well (384-well plate format) on the number of GFP-BICD2N-FRB and PTS-RFP-FKBP spots. Cells were seeded for 72 h prior to treatment with rapamycin (2 nM). Datapoints represent mean aggregation at well level (minimum of three wells analysed per condition). Error bars, S.D.. 1500 cells per well were seeded for the genome-wide screen.

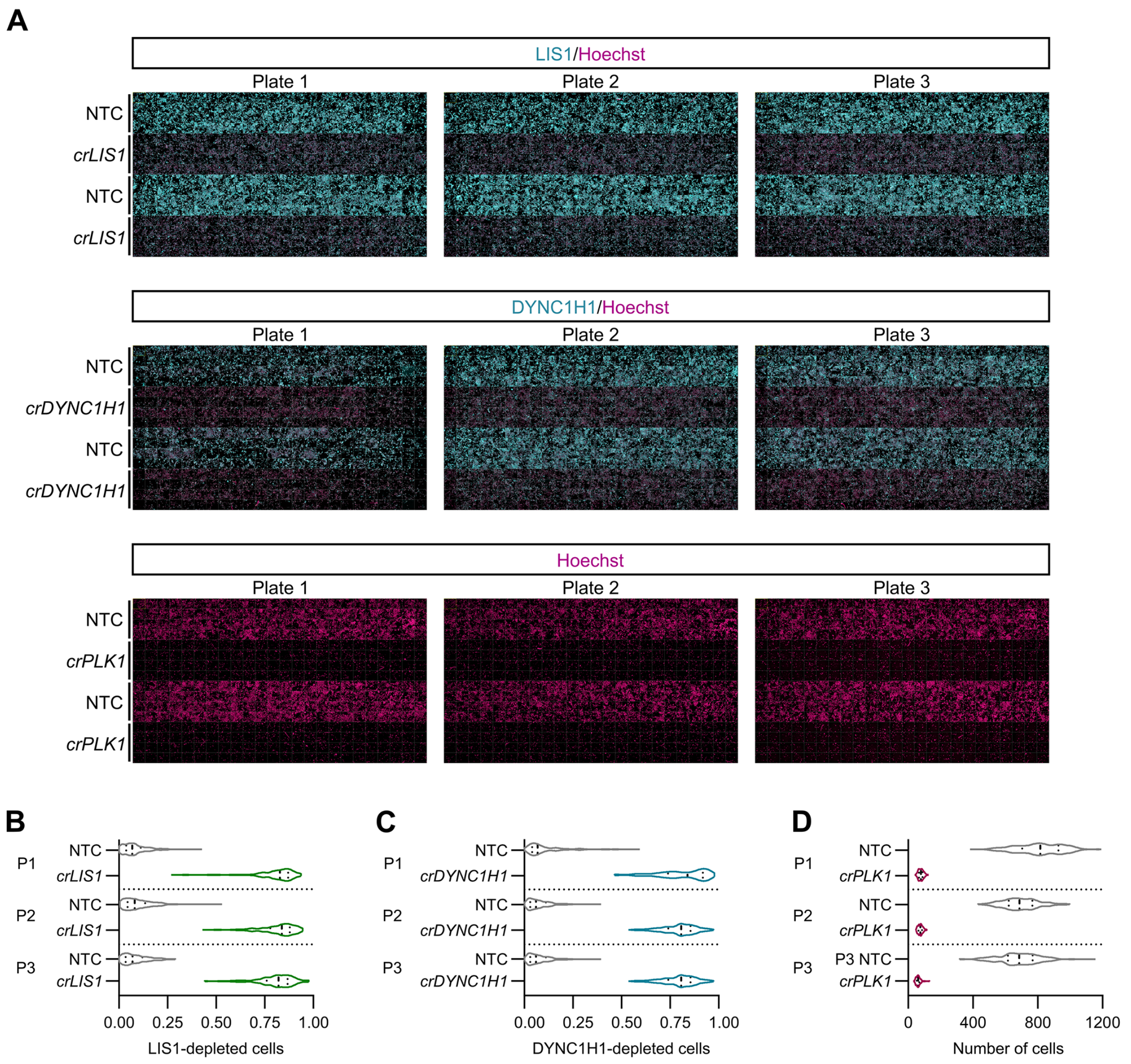


**Supplementary figure 4. Assay scaling for high-throughput editing. A)** Representative low magnification view of 384-well plate regions showing consistent editing in U-2 OS PEX cells treated with *crLIS1*, *crDYNC1H1*, and *crPLK1*. *crLIS1* and *crDYNC1H1* activity was assessed by immunostaining for the target proteins, whereas activity of *crPLK1* was read out by a reduction in cell number (revealed by Hoechst staining). **B – D)** Violin plots (median, bold line; first/third quartile, dashed lines) of frequency of cells depleted for LIS1 (B) or DYNC1H1 (C), or the number of cells (D), after transfection with *crLIS1*, *crDYNC1H1* or *crPLK1*, respectively. Datapoints represent mean aggregation at the well level (minimum of 100 cells from at least four wells analysed) for three individual plates (P).

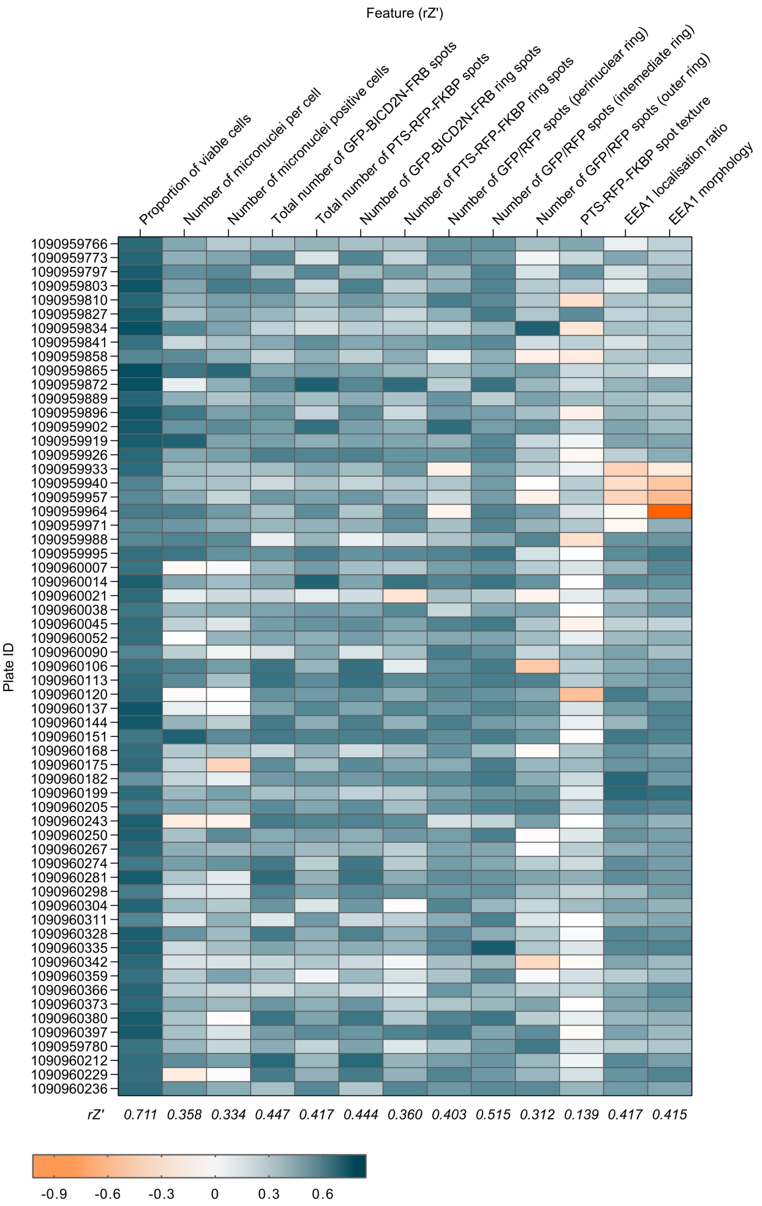


**Supplementary figure 5. Quality control of assay endpoints used in the genome-wide screen.** Heatmap displaying rZ′ score of features used for hit calling from the screen data. All endpoints were normalised by NTC vs *crLIS1* except for cell number, which was normalised by NTC vs *crPLK1*. The displayed rZ' scores represent the corrected rZ' for the individual features for the entire screening set. See Supplementary table 1 for the complete dataset.

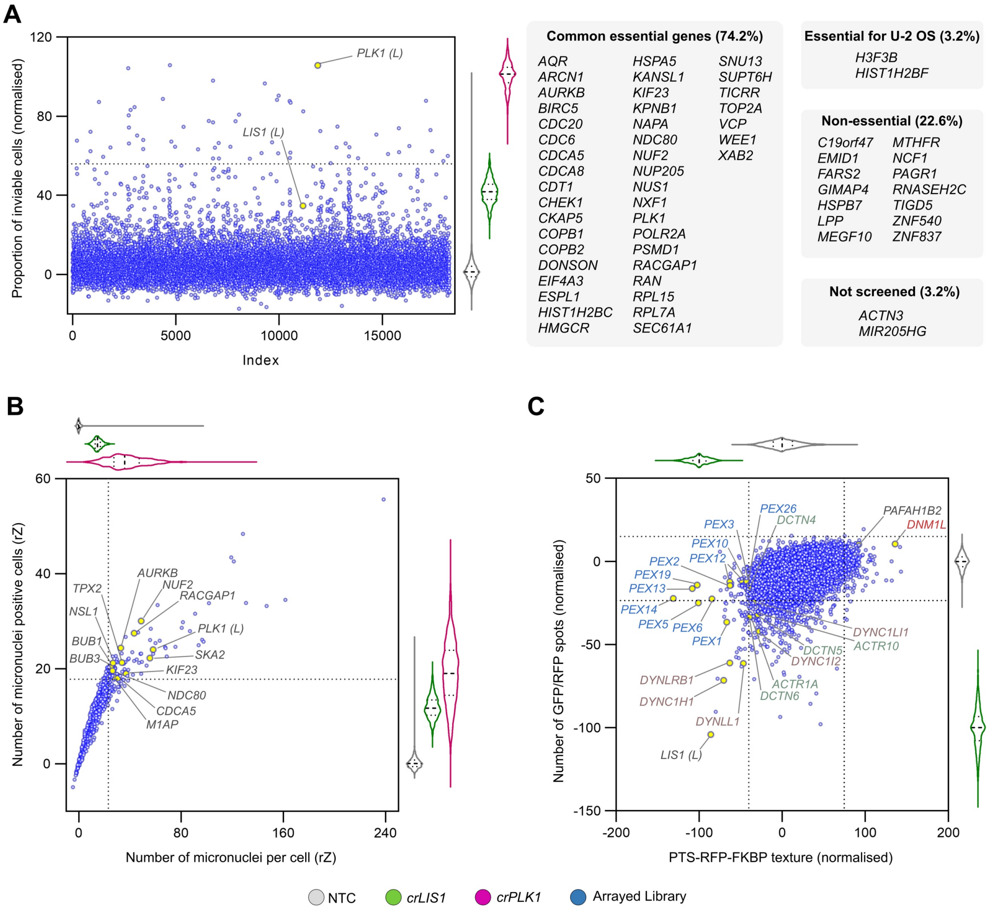


**Supplementary figure 6. Additional genome-wide screen endpoints. A)** Effects of arrayed library crRNAs on cell viability and comparison to results from previous cell viability studies. Scatter plot of library results and corresponding violin plot of controls (median, bold dashed line; first/third quartile, dashed lines; colour code in key at bottom of figure) of proportion of inviable cells (gated based on nuclear morphology of NTC cells). Data points represent normalised values based on the neutral control (NTC, 0) and lethal editing control (*crPLK1*, 100). Dashed line on the y-axis represents 2.5*S.D. of *crLIS1*, the threshold for calling lethal crRNAs. Library copies of *crPLK1* and *crLIS1* are labelled with (L). The genes targeted by the lethal crRNAs were cross-referenced with their gene essentiality categorisation from the Cancer Dependency Map project (DepMap 22Q2 Public+Score; https://depmap.org/portal): ‘common essential genes’ are classed as essential for growth and survival in at least 90% of cancer cell lines; ‘essential for U-2 OS’ genes are those classed as not essential across multiple cell lines but essential in U-2 OS cells; ‘non-essential’ genes are those not identified as essential in a panel of sensitive cell lines; ‘not screened’ genes are those that are not represented in the DepMap screening dataset. Numbers in parentheses refer to percentage of essential genes in our dataset that were found in each Depmap category. **B)** Effects of arrayed library crRNAs on micronuclei incidence. Scatter plot of library results and corresponding violin plot of controls (median, bold dashed line; first/third quartile, dashed lines; colour code in key at bottom of figure) for number of micronuclei per cell (x-axis) and number of cells containing micronuclei (y-axis). Data points represent rZ normalisation (central reference = NTC). Dashed lines on the x- and y-axes represent 2.5*S.D. of *crLIS1*, the threshold for calling crRNAs that cause micronuclei abnormalities. Library copy of *crPLK1* is labelled with (L). Labelled genes were functionally enriched (FDR ≤ 0.005) for gene ontology terms associated with regulation of chromosome segregation, including ‘nuclear division’, ‘mitotic sister chromatid segregation’ and ‘nuclear chromosome segregation’ (http://bioinformatics.sdstate.edu/go/). **C)** Effects of arrayed library crRNAs on PTS-RFP-FKBP texture and number of PTS-RFP-FKBP and GFP-BICD2N-FRB spots. Scatterplot of library results and corresponding violin plots of controls (median, bold dashed line; first/third quartile, dashed lines; colour code in key at bottom of figure) of normalised values based on the neutral control (NTC, 0) and inhibition (*crLIS1*, -100). Dashed lines on both axes represent ± 2.5*S.D. of NTC and arrayed library, the threshold for hit calling. Both endpoints were generated from a linear discriminant analysis; PTS-RFP-FKBP texture was generated from six textural features based on filtered images, while ‘GFP/RFP spots’ was generated by two features (total number of GFP-BICD2N-FRB and PTS-RFP-FKBP spots). Core components of the dynein complex and dynactin complex that met the threshold for hit calling for either endpoint are labelled in purple and teal text, respectively. Library copy of *crLIS1* is labelled with (L). Known peroxisome biogenesis genes are labelled in blue text (note that there is a duplicate of the *PEX12* crRNA pool in the library). *crPAFAH1B2* and *crDNM1L* are labelled as examples of crRNAs that affect PTS-RFP-FKBP texture in the opposite way to crRNAs against dynein-dynactin components.

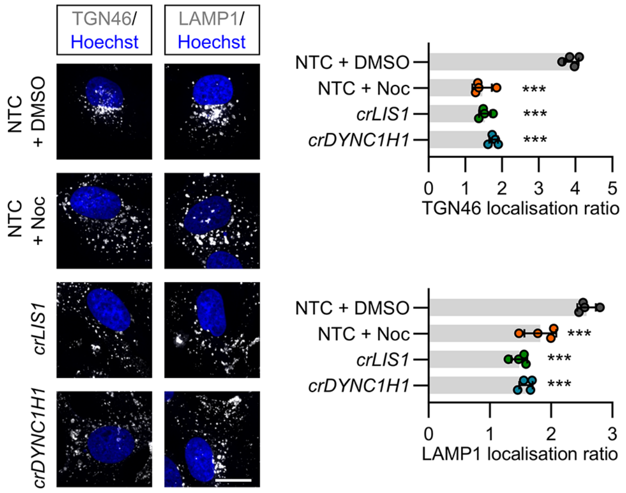


**Supplementary figure 7. Confirmation that dynein promotes perinuclear enrichment of Golgi and lysosomal membranes in U-2 OS cells.** Representative images and quantification of dispersion of *trans*-Golgi network (TGN46) and lysosomal membranes (LAMP1) in U-2 OS cells following the indicated treatments (Noc, nocodazole). Bar graphs show ratio of spot number in the perinuclear region vs the peripheral region (lower values indicate increased dispersion). Data points represent mean aggregation at the well level (minimum of 100 cells analysed per well, four wells analysed per condition). Error bars signify S.D.. ****p*<0.001 (one-way ANOVA with Dunnett's multiple comparison vs NTC +DMSO). Scale bar, 20 µm.

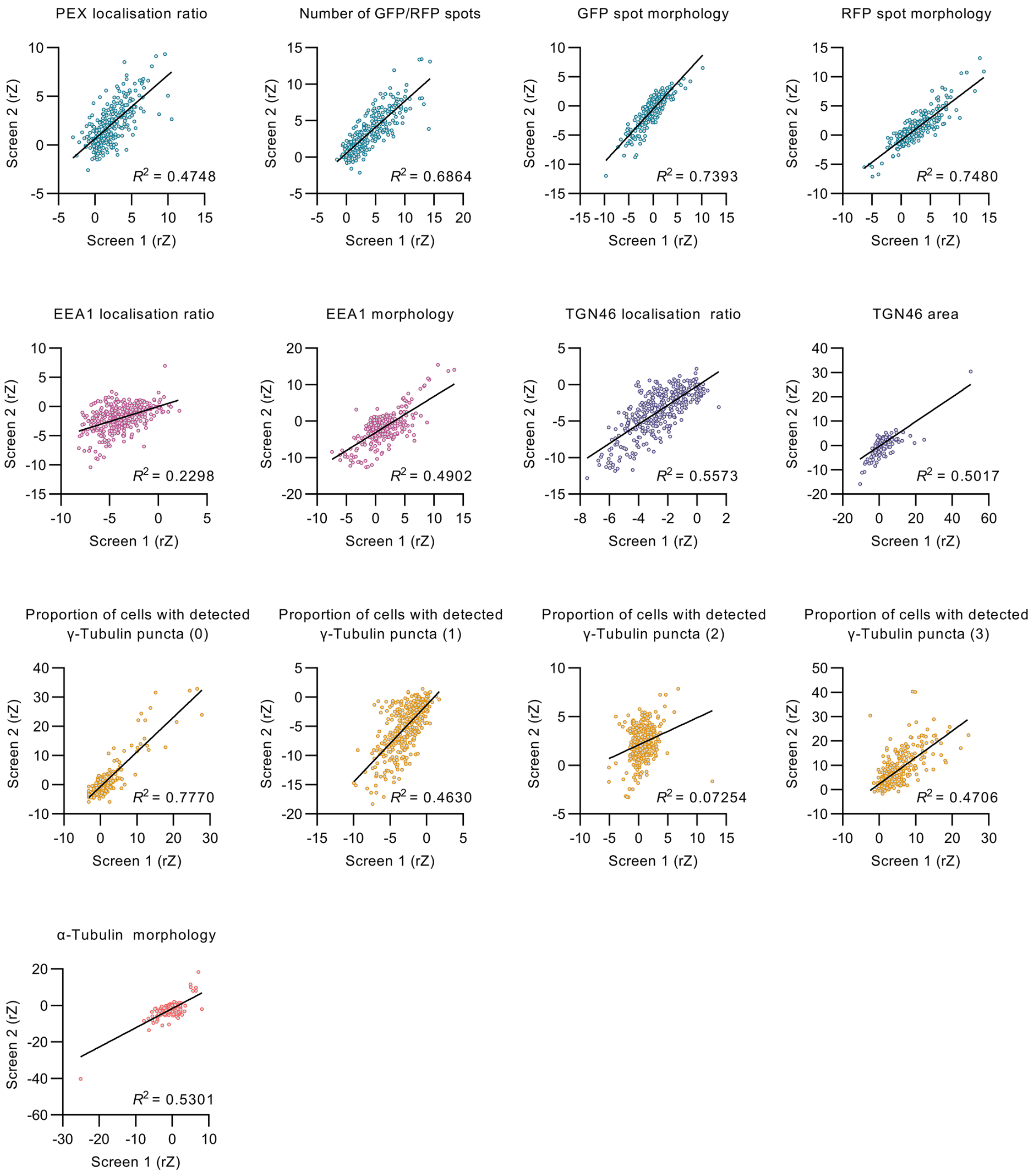


**Supplementary figure 8. Reproducibility of secondary screen runs.** Linear regression was performed on the main endpoints used in the two independent runs in the secondary screen. Data points represent the median rZ (central reference = NTC) of individual crRNA pools. The only metric with a poor $R^2$ score (proportion of cells with two γ-Tubulin puncta) was not used for hit calling.

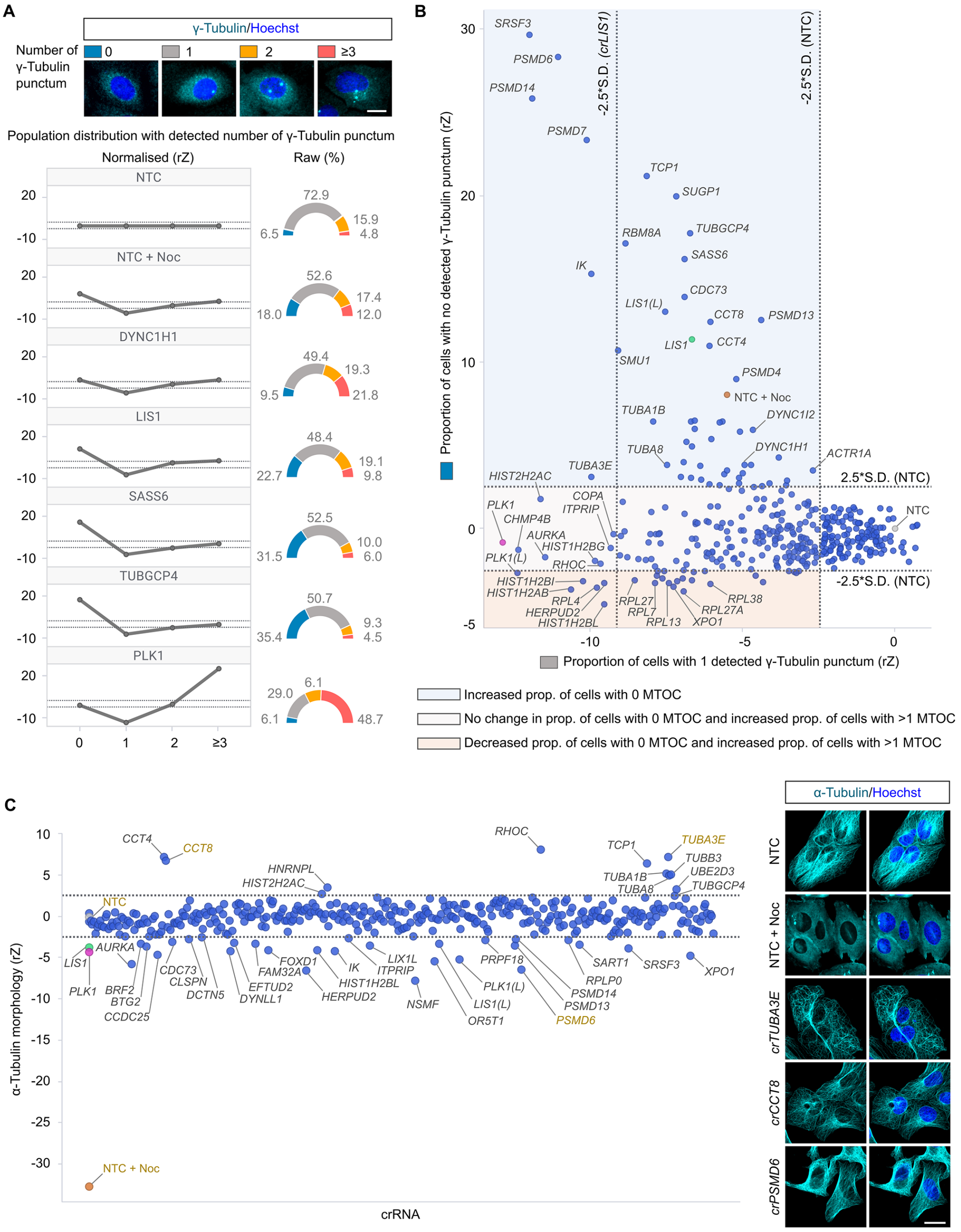


**Supplementary figure 9. Analysis of γ-Tubulin and α-Tubulin signals in the secondary screen. A)** Representative images of unmodified U-2 OS cells with different numbers of detected γ-Tubulin puncta (from population of NTC controls) and distribution of γ-Tubulin puncta numbers in cells treated with nocodazole (Noc) or the indicated crRNA pools. Line charts show normalised data (rZ normalisation, central reference = NTC) and pie charts show raw data (percentage of cells). Scale bar, 50 µm. **B)** Scatterplot of normalised values (rZ normalisation, central reference = NTC) for proportion of cells with one versus no detected γ-Tubulin punctum. crRNAs with values to the left of the vertical -2.5*S.D. (NTC) cut-off line were classed as causing a decrease in the proportion (prop.) of cells with one detected γ-Tubulin punctum (243/377 crRNA pools). This phenotype could be associated with either an increase in the proportion of cells with more than one MTOC (193 crRNAs (values below the horizontal 2.5*S.D. (NTC) cut-off line; regions shaded in pink and dark pink)) or an increase in the proportion of cells with no MTOC (50 crRNAs (values above the horizontal 2.5*S.D. (NTC) cut-off line; region shaded in blue)). An additional threshold based on *crLIS1* controls (-2.5* S.D.) was introduced for the one punctum feature to categorise crRNAs that yielded a stronger phenotype than *crLIS1*. **C)** Quantification of α-Tubulin morphology in U-2 OS PEX cells generated by linear discriminant analysis (rZ, central reference = NTC) and representative immunofluorescence images of examples (labelled in gold text in the plot). In B and C, library copies of *crPLK1* and *crLIS1* are labelled with (L). Scale bar, 25 µm. Data points in A – C represent mean values of two independent experiments with at least four wells per crRNA per experiment. Hits were defined as exceeding ± 2.5*S.D. of (A-C) NTC and/or (B) *crLIS1*. See Supplementary table 6 for full dataset.

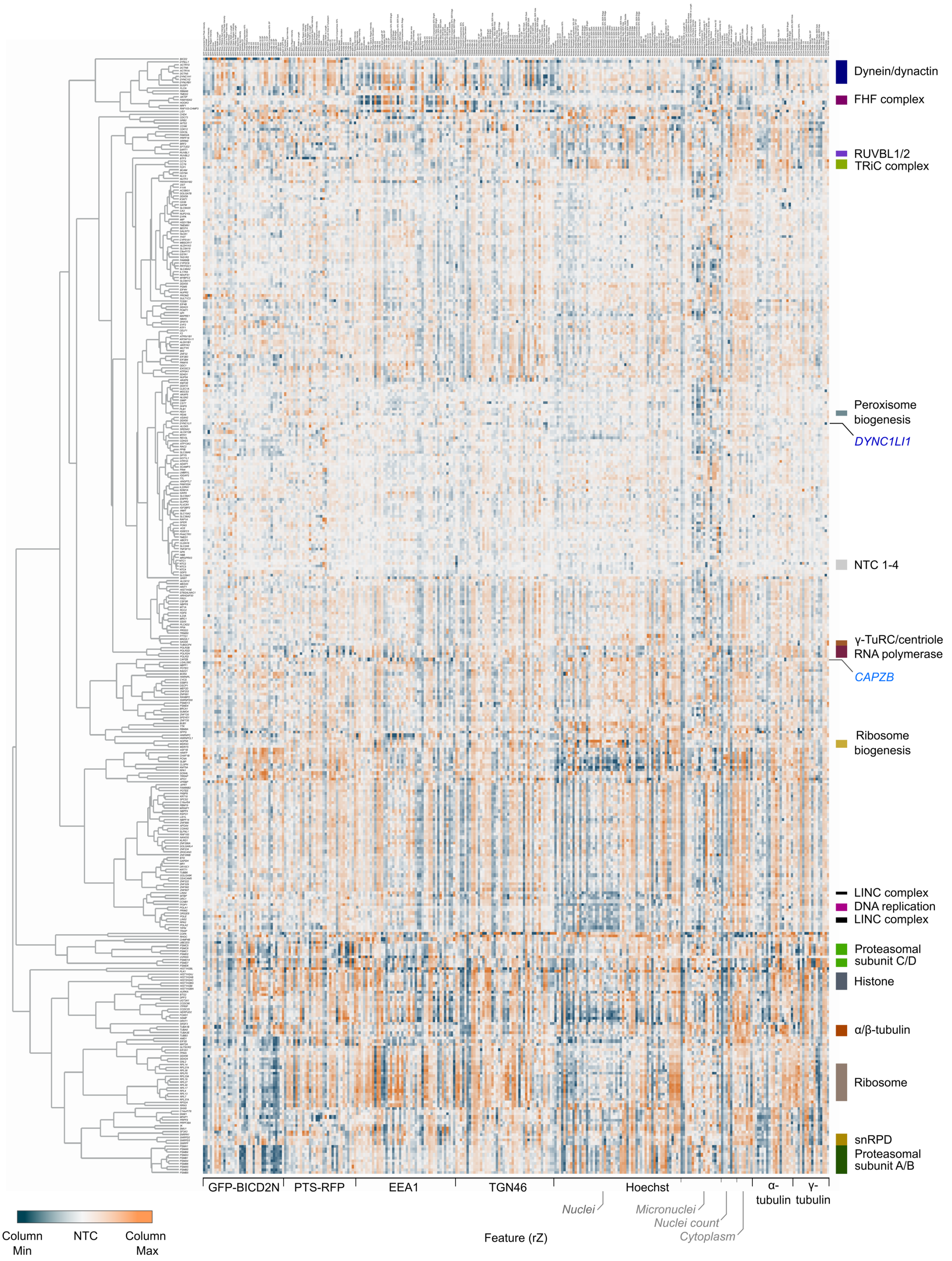


**Supplementary figure 10. Overview of functional clusters from image-based profiling of secondary screen data.** Displayed is the phenotypic feature heatmap generated by hierarchical clustering. The data, including gene names and feature titles, can be explored by zooming in within the additional file 'Supplementary phenotypic heatmap', as well as in Supplementary table 7. The scale of rZ values (central reference = NTC) was adjusted based on the min and max values of individual features. Labels to the right highlight a subset of functional clusters, as well as the positions of the four NTC (neutral control) crRNAs and crRNAs targeting *CAPZB* and *DYNC1LI1*. 'Cytoplasm' refers to features associated with the background Hoescht staining in the cytoplasm.

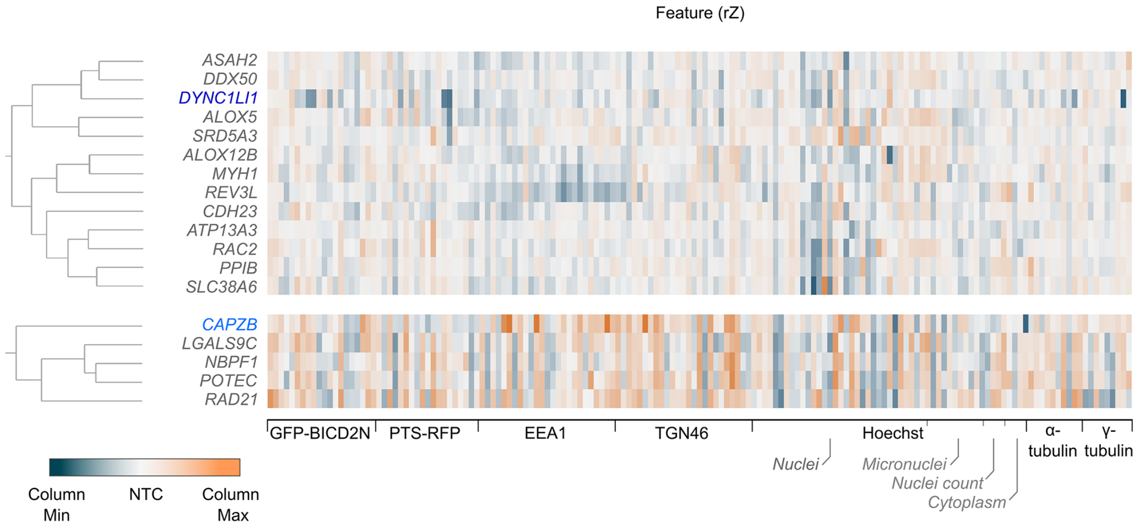


**Supplementary figure 11. Phenotypic feature heatmaps for *CAPZB*- and *DYNC1LI1*-containing clusters.** The scale of rZ values (central reference = NTC) is adjusted based on the min and max values of individual features. ‘Cytoplasm’ refers to features associated with the background Hoescht staining in the cytoplasm.

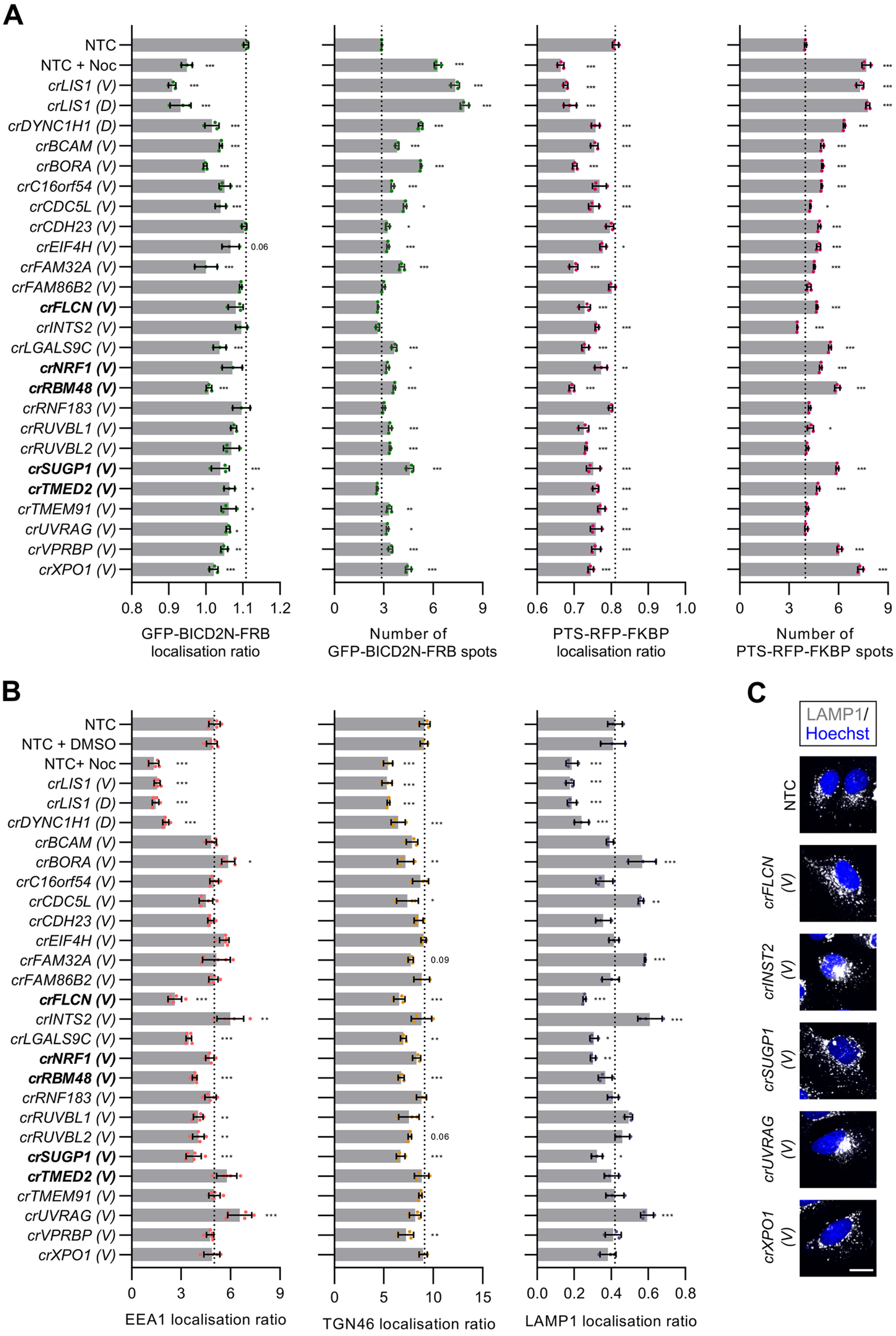


**Supplementary figure 12. Readouts of the orthogonal cargo localisation screen. A, B)** Bar graphs showing readouts generated from the screen with VBC-generated crRNAs. Quantification of (A) localisation ratio (perinuclear vs peripheral) and total number for GFP-BICD2N-FRB and PTS-RFP-FKBP spots in U-2 OS PEX cells treated with rapamycin and (B) localisation ratio (perinuclear vs peripheral) of EEA1, TGN46 or LAMP1 spots in unmodified U-2 OS cells. '(V)' and '(D)' indicate crRNAs synthesised based on the VBC score or from the original Discovery set, respectively. Data points represent mean aggregation from at least three independent experiments (minimum of 100 cells analysed per well, four wells analysed per condition). EEA1 ratio values were log-transformed for normal distribution. Bold lettering indicates crRNAs that were novel components of the 'dynein-dynactin' cluster of phenotypic profiles. Error bars signify S.D.. **p*<0.05, ***p*<0.01, ****p*<0.001 (one-way ANOVA with Dunnett's multiple comparison against NTC). **C)** Representative images of LAMP1 staining in U-2 OS cells treated with the indicated crRNAs. Scale bar, 20 µm.

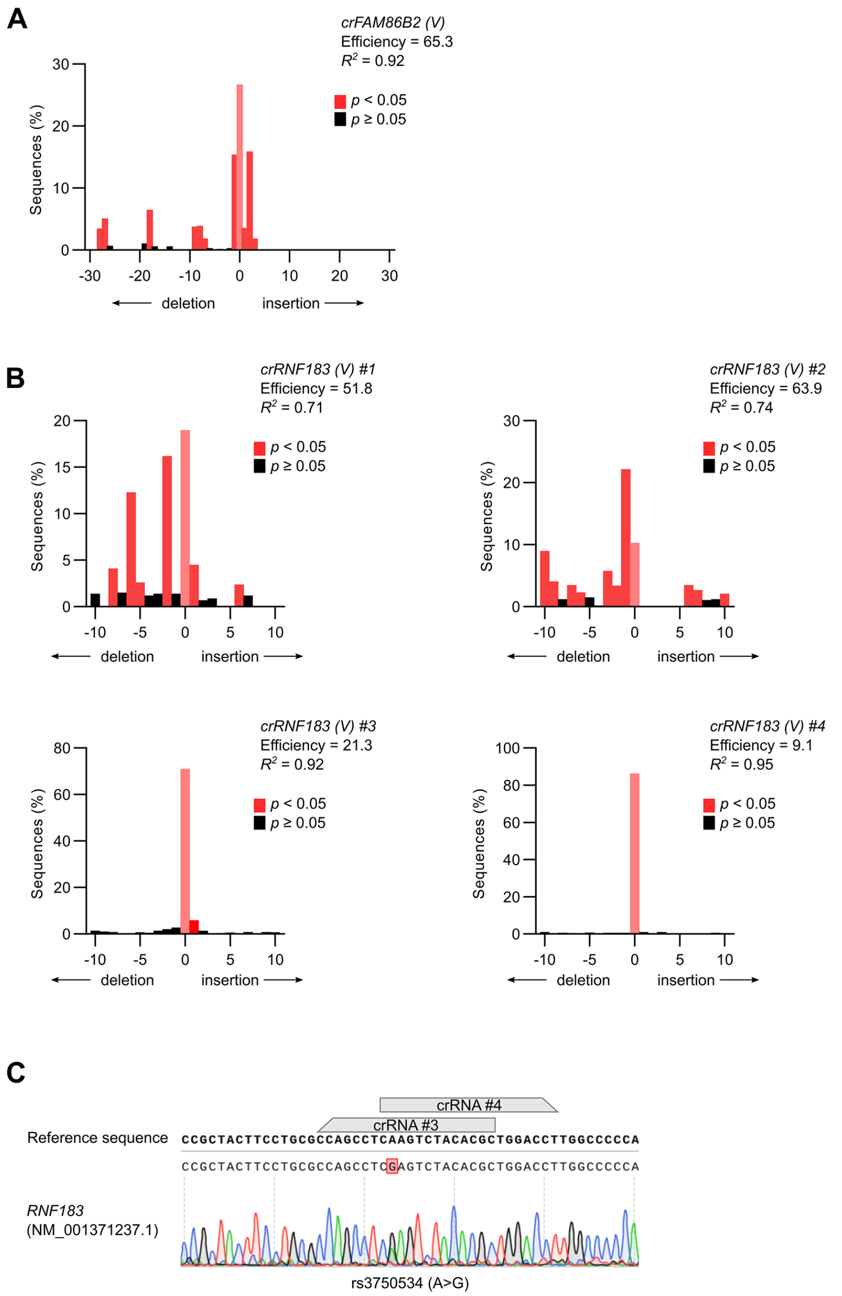


**Supplementary figure 13. TIDE analysis for VBC-derived *FAM86B2* and *RNF183* crRNAs. A, B)** Indel distribution derived from sequences of target regions in unmodified U-2 OS cells transfected with VBC-derived *FAM86B2* (A) or *RNF183* (B) crRNAs. Corresponding sequences from cells transfected with NTC crRNAs were used as a reference. Insufficient targeting of *crFAM86B2 pool* may be due to all four crRNAs within the pool targeting overlapping regions, and therefore competing with each other. Efficiency scores are out of 100. **C)** Incomplete targeting with the *crRNF183* pool in U-2 OS cells may be associated with an overlapping target region and/or spanning of a common polymorphism (rs3750534) within a target sequence for *crRNF183* (V) #3 and *crRNF183* (V) #4, which have particularly low activity.

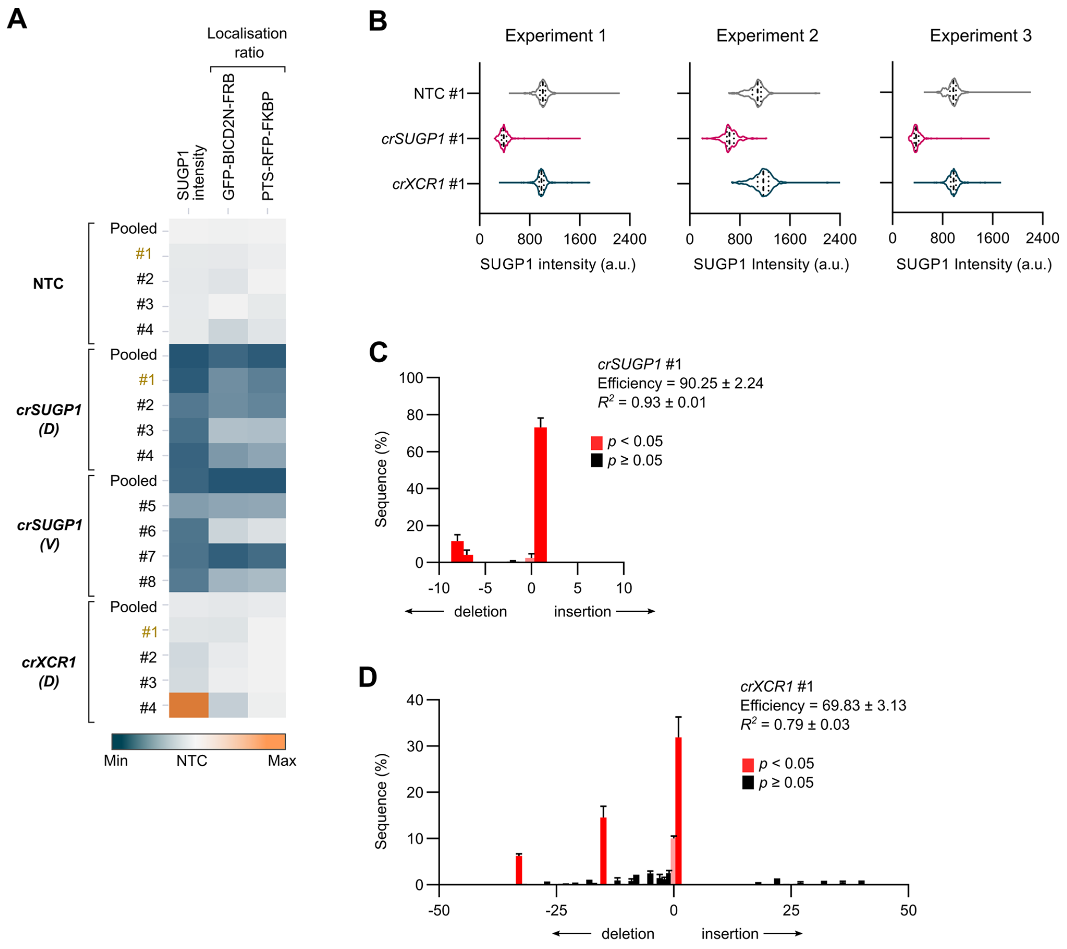


**Supplementary figure 14. Guide validation and phenotyping for *crSUGP1*. A)** Phenotyping of individual and pooled *SUGP1* crRNAs. Heatmap displaying quantification of SUGP1 protein signal and localisation ratio (perinuclear vs peripheral) of GFP-BICD2N-FRB and PTS-RFP-FKBP spots in U-2 OS PEX cells with the indicated treatments. ‘D’ and ‘V’ indicates source of crRNA design (D, Discovery; V, VBC score). Data points represent mean aggregation of four wells, with a minimum of 100 cells analysed per conditiion. Colour scale of individual features was adjusted based on their min and max raw values. The guides selected for RNA-seq are labelled in gold text. **B – D)** Quality control for RNA samples submitted for RNA-seq. (B) Violin plots of intensity of SUGP1 protein signal at the single cell level (median, bold line; first/third quartile, dashed lines; minimum of 100 cells analysed per condition). (C, D) Indel distribution derived from sequences of crRNA target region of U-2 OS cells transfected with *crSUGP1* #1 (C) or *crXCR1* #1 (D). Sequences from cells transfected with the NTC #1 crRNA were used as the reference control. Bar graphs display mean ± S.D.. Efficiency score is out of 100.

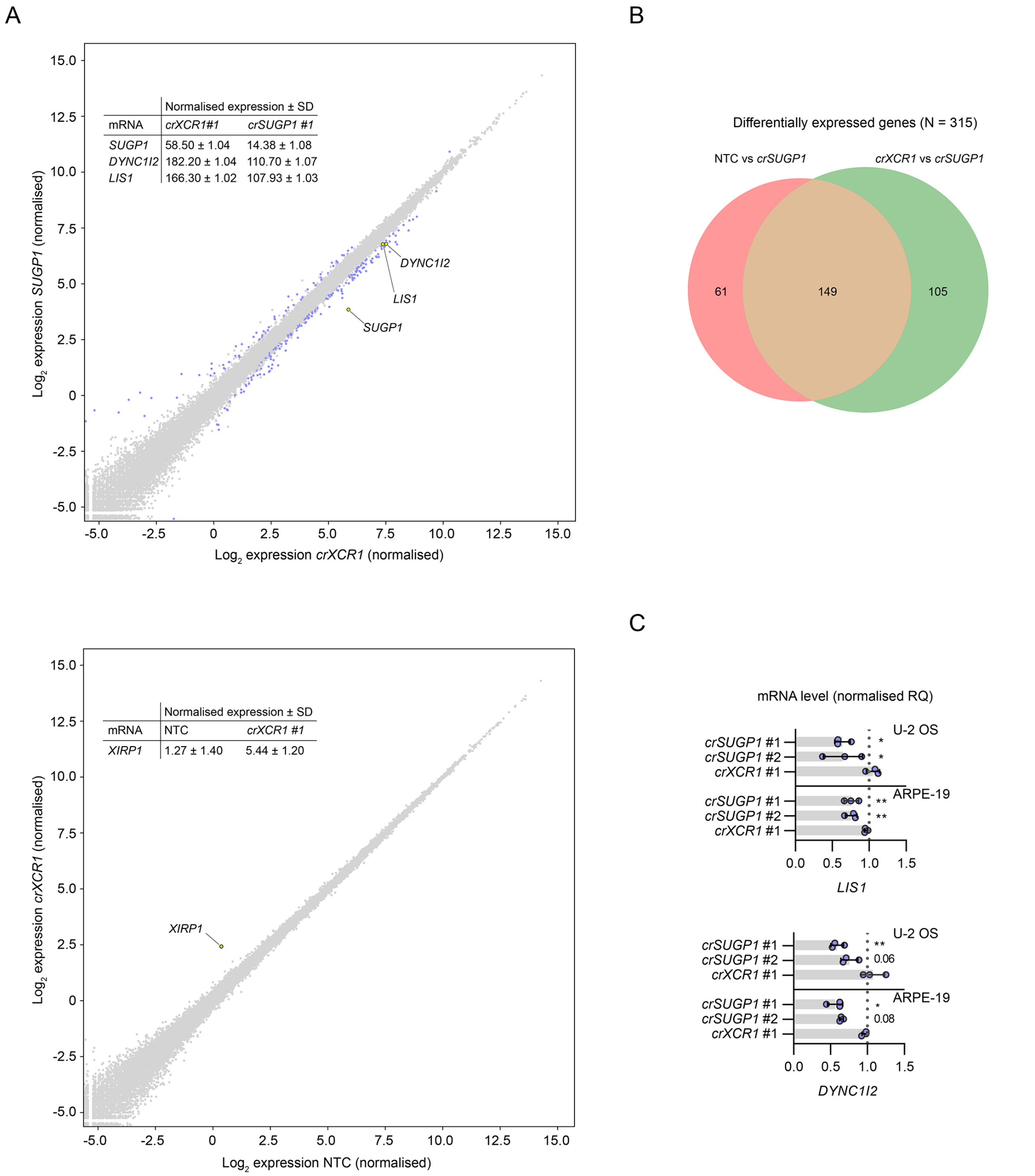


**Supplementary figure 15. Supplementary data for differential expression analysis. A)** Scatter plot of mRNA abundance for (top panel) *XCR1*-edited vs *SUGP1*-edited U-2 OS cells and (bottom panel) NTC vs *XCR1*-edited U-2 OS cells (mean $\log_2$ normalised values from three independently performed experiments). mRNAs meeting the threshold for inclusion (minimum absolute $\log_2$ normalised fold change ≥ 0.5 and FDR ≤ 0.05) are labelled in blue, except (top panel) *SUGP1*, *DYNC1I2* and *LIS1* and (bottom panel) *XIRP1* (the only differentially expressed gene in the NTC vs *crXCR1* comparison), which are labelled in yellow. Inset tables show non-logarithmic values for (top panel) *SUGP1*, *DYNC1I2* and *LIS1* and (bottom panel) *XIRP1* mRNAs. See Supplementary table 8 for full results. **B)** Venn diagram showing overlap of differentially expressed genes in the NTC *vs crSUGP1* and *crXCR1* vs *crSUGP1* comparisons. **C)** Quantification of *LIS1* and *DYNC1I2* mRNA level, determined by TaqMan-based real-time quantitative PCR, in *SUGP1*-edited and *XCR1*-edited U-2 OS and ARPE-19 cells. Data points represent mean of three independent experiments (RQ = relative quantification based on NTC). Error bars signify S.D.. **p*<0.05, ***p*<0.01 (one-way ANOVA with Dunnett's multiple comparison against NTC).

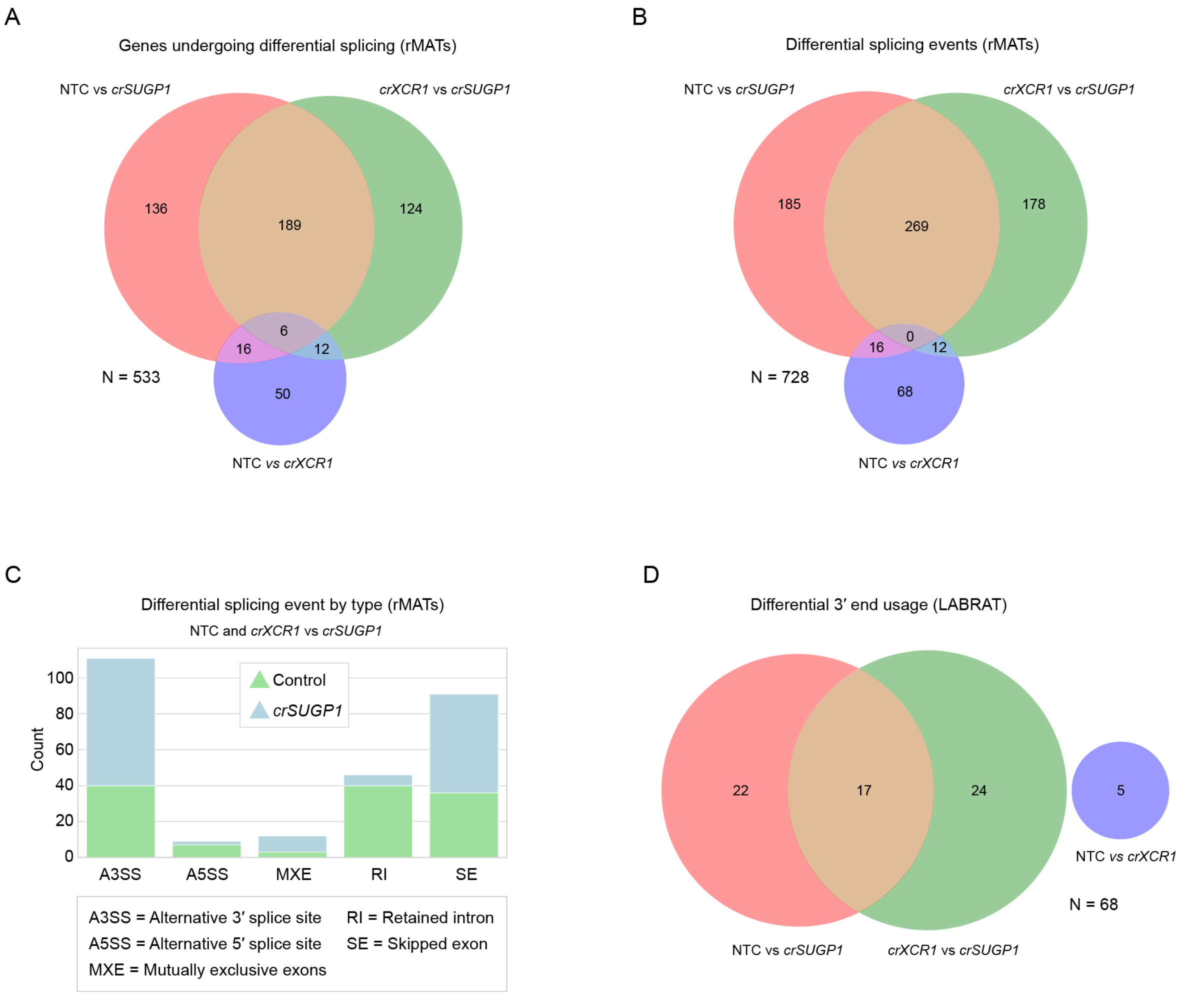


**Supplementary figure 16. Supplementary data for differential splicing analysis. A, B)** Venn diagrams showing overlap of genes that undergo differential splicing (A) and differential splicing events (B) in the datasets, as determined with rMATs (note that some genes have more than one differential splicing event). Threshold for classifying an event as differential was: absolute IncLevelDifference ≥ 0.2, total read count (inclusion count + skipping count) ≥ 10 and FDR ≤ 0.05. See Supplementary table 9 for full results. **C)** Classes of alternative splicing events common to both comparisons (i.e. NTC vs *crSUGP1* or *crXCR1* vs *crSUGP1*), as identified by rMATS. Blue and green represent events that were enriched in control and *crSUGP1* samples, respectively. rMATS reports five splicing categories: i) alternative 3’ splice sites (A3SS); ii) alternative 5’ splice sites (A5SS); iii) mutually exclusive exons (MXE); iv) retained introns (RI), and v) skipped exons (SE). The A3SS and A5SS events involve the splicing together of two exons separated by a single intron. For A3SS events, alternative splicing causes a downstream exon to extend partially into neighbouring intronic sequence. A5SS is defined by an alternative splicing event causing an upstream exon to extend partially into the adjoining intron. Mutually exclusive exons describe the splicing of adjacent exons (separated by a single intron) in which one exon is retained but the other is excluded, or vice versa. The graph reports MXE events in which the upstream exon was selected. Events classified as RI are those in which an intron is not spliced out and hence is retained in the mature transcript. SE denotes splicing events in which an exon is skipped over and not included in the processed RNA molecule. **D)** Venn diagrams showing overlap of differential 3′-end usage events in the datasets, as determined by LABRAT. LABRAT quantifies alternative polyadenylation (APA) sites and reports upstream or downstream shifts in the usage of those sites for each gene as compared to the control (see Supplementary table 10 for full results). The threshold for classifying an event as differential was $\Delta\psi \geq 0.05$ and FDR ≤ 0.05.

### SUPPLEMENTARY TABLE LEGENDS

**Supplementary table 1. Features and data acquired in the genome-wide screen (Excel file)**

List of acquired features (Tab 1), full dataset (Tab 2) and rZ' scores for features used for hit calling (Tab 3). The annotated gene ID is followed by Genecard ID, with the well type indicated (Neutral control, NTC; Editing control, *crPLK1*; Inhibitor control, *crLIS1*; Library, arrayed library). Note that the *LIS1* library copy is referred to by the official Genecard ID: *PAFAH1B1*. Datapoints represent means aggregated from four fields of view per sample. Individual features are indicated as either raw (unprocessed) or in normalised form as appropriate (normP, normalised against *crPLK1*; normL, normalised against *crLIS1*; rZ, NTC as central reference). Wells with no data or incomplete data for certain features may indicate a failure in imaging, insufficient cells (*e.g.* lethal crRNAs) and/or low quality samples that resulted in insufficient raw data for analysis. Wells with RNA dispensing failure are indicated, with control wells masked for normalisations. $\alpha$-Tubulin data are included for completeness but values were too variable across the plates to draw conclusions about effects of crRNA pools on the microtubule network. In Tab 3, all rZ' scores were calculated via normalisation with NTC vs *crLIS1* except for 'proportion of viable cells', which was normalised by NTC vs *crPLK1*.

**Supplementary table 2. Lists of hits identified using different features of cellular organisation (Excel file)**

List of features from the genome-wide screen used for hit calling, with the indicated normalised data (normP, normalised against *crPLK1*; normL, normalised against *crLIS1*; rZ, NTC as central reference). The selected endpoints include those related to cell viability, nucleus morphology, peroxisome relocalisation assay and EEA1 morphology. See Supplementary table 1 for the complete dataset.

**Supplementary table 3. Components of the dynein machinery identified in assay endpoints (Excel file)**

Summary of dynein-dynactin components and known co-factors and the hit criteria they met in the genome-wide screen. All of the listed components are associated with a dispersion phenotype, except for *crPAFAH1B2*, which was associated with increased clustering of EEA1.

**Supplementary table 4. List of genes taken forward to secondary screens (Excel file)**

**Supplementary table 5. K-means clusters and cargo localisation scores from secondary screen (Excel file)**

Cargo localisation ratio (PEX, EEA1 and TGN46) of the individual crRNAs categorised by clusters identified by K-means. Values represent rZ normalisation (central reference = NTC).

**Supplementary table 6. γ-Tubulin and $\alpha$-Tubulin datasets from secondary screen (Excel file)**

Analysis of $\alpha$-Tubulin morphology and γ-Tubulin spot number (population distribution with detected number of puncta – 0,1,2,3). Values represent rZ normalisation (central reference = NTC).

**Supplementary table 7. Phenotypic fingerprints for clustering (Excel file)**

Non-redundant features (n = 278) and the respective scores for crRNAs subjected to hierarchical clustering. Values represent rZ normalisation (central reference = NTC). The annotated gene ID is followed by Genecard ID. The display order of features (columns) and crRNAs (rows) in this table is arranged as in Supplementary figure 10 and the Supplementary phenotypic heatmap file.

**Supplementary table 8. Differentially expressed genes in NTC, *crSUGP1* and *crXCR1* samples (DEseq) (Excel file)**

In the last three columns, TRUE and FALSE refers to, respectively, the presence and absence of statistically significant differences for the indicated comparisons.

**Supplementary table 9. Differential slicing events in NTC, *crSUGP1* and *crXCR1* samples (rMATS) (Excel file)**

In the last three columns, TRUE and FALSE refers to, respectively, the presence and absence of statistically significant differences in alternative splicing (AS) for the indicated comparisons. Further details of specific parameters can be found in ref. 79[79].

**Supplementary table 10. Differential 3′-end usage events in NTC, *crSUGP1* and *crXCR1* samples (LABRAT) (Excel file)**

Negative and positive delta ψ values refer, respectively, to downstream and upstream shifts in APA usage in the *crSUGP1* samples. APA sites can be contained within the same exon ('tandem UTR' (TUTR) genes) or different exons ('alternative last exon'; ALE genes). Some genes may constitute a mixture of these two categories, *i.e.* possessing APA sites that co-localise to the same exon as well as having APA sites in other exons; such genes are termed 'mixed'. In the last three columns, TRUE and FALSE refers to, respectively, the presence and absence of statistically significant differences for the indicated comparisons. Only genes that have significant FDR values (≤ 0.05) are shown. Δψ for genes that did not meet this threshold are indicated with #NA. Further details can be found in ref. 80.

**Supplementary table 11. Primary antibodies used for immunofluorescence (Word file)**

**Supplementary table 12. Secondary antibodies used for immunofluorescence (Word file)**
