## Supplementary figures and images for "Genome-scale requirements for dynein-based trafficking revealed by a high-content arrayed CRISPR screen"

### Supplementary phenotypic heatmap

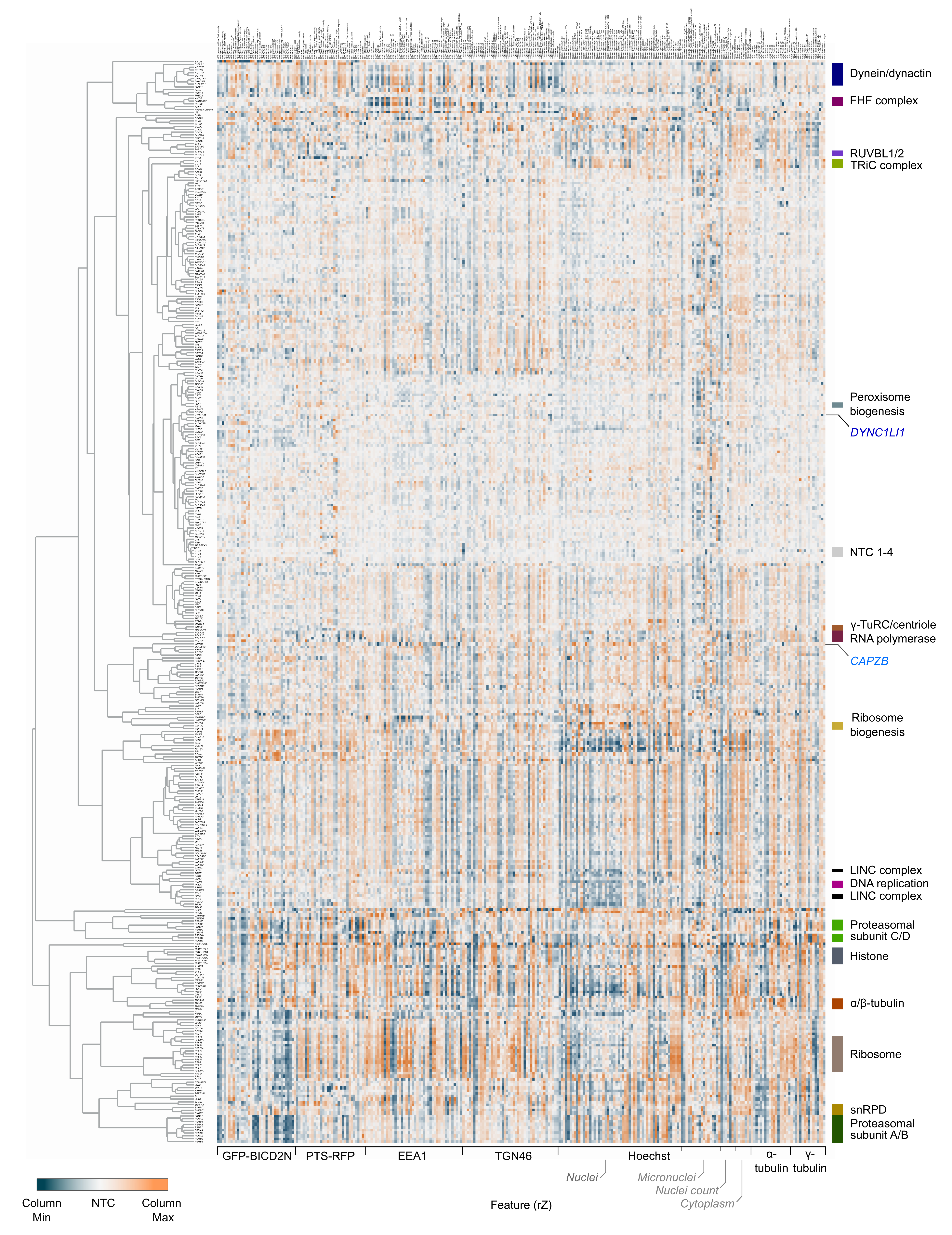
