## Supplementary table 11 for "Genome-scale requirements for dynein-based trafficking revealed by a high-content arrayed CRISPR screen"

### Supplementary table 11. Primary antibodies used for immunofluorescence

| Antibody target | Host | Catalog number | Vendor | Dilution |
|---|---|---|---|---|
| HA | Rabbit | 3724 | Cell Signaling Technology | 1:500 |
| HA | Mouse | 2367 | Cell Signaling Technology | 1:500 |
| α-Tubulin | Mouse | T9026 | Sigma Aldrich | 1:3000 |
| α-Tubulin | Rat | MCA77G | Bio-Rad | 1:3000 |
| LIS1 | Mouse | H00005048-M03 | Novus | 1:1000 |
| DYNC1H1 | Rabbit | 12345-1-AP | Proteintech | 1:1000 |
| DYNC1I2 | Rabbit | HPA053987 | Atlas | 1:500 |
| EEA1 | Rabbit | 3288 | Cell Signaling Technology | 1:3000 |
| TGN46 | Sheep | AHP500G | Bio-Rad | 1:3000 |
| γ-Tubulin | Mouse | T5326 | Sigma Aldrich | 1:1000 |
| LAMP1 | Rabbit | 9091 | Cell Signaling Technology | 1:3000 |
| SUGP1 | Rabbit | HPA004890 | Atlas | 1:1000 |
| PPIB | Rabbit | ab16045 | Abcam | 1:1000 |
| DCTN1 | Mouse | 612708 | BD | 1:1000 |
| GOLGA3 | Rabbit | 21193-1-AP | Proteintech | 1:500 |
| TARDBP | Rabbit | 12892-1-AP | Proteintech | 1:1000 |
| V5 | Rabbit | 13202 | Cell Signaling Technology | 1:1000 |
| V5-AF647 | Mouse | 451098 | Thermo Fisher | 1:1000 |
| FLAG | Mouse | TA50011-100 | Origene | 1:1000 |
