## Supplementary table 12 for "Genome-scale requirements for dynein-based trafficking revealed by a high-content arrayed CRISPR screen"

**Supplementary table 12. Secondary antibodies used for immunofluorescence**

| Antibody | Catalog number | Vendor | Dilution |
|---|---|---|---|
| Donkey anti-Mouse IgG (H+L) Highly Cross-Adsorbed Secondary Antibody, Alexa Fluor 488 | A-21202 | Thermo Fisher | 1:500 |
| Donkey anti-Rabbit IgG (H+L) Highly Cross-Adsorbed Secondary Antibody, Alexa Fluor 488 | A-21206 | Thermo Fisher | 1:500 |
| Goat anti-Mouse IgG (H+L) Highly Cross-Adsorbed Secondary Antibody, Alexa Fluor 568 | A-11031 | Thermo Fisher | 1:500 |
| Donkey anti-Rabbit IgG (H+L) Highly Cross-Adsorbed Secondary Antibody, Alexa Fluor 568 | A10042 | Thermo Fisher | 1:500 |
| Donkey anti-Mouse IgG (H+L) Highly Cross-Adsorbed Secondary Antibody, Alexa Fluor 647 | A-31571 | Thermo Fisher | 1:500 |
| Donkey anti-Rabbit IgG (H+L) Highly Cross-Adsorbed Secondary Antibody, Alexa Fluor 647 | A-31573 | Thermo Fisher | 1:500 |
| Donkey anti-Rat IgG (H+L) Highly Cross-Adsorbed Secondary Antibody, Alexa Fluor 647 | A78947 | Thermo Fisher | 1:500 |
| Donkey anti-Sheep IgG (H+L) Cross-Adsorbed Secondary Antibody, Alexa Fluor 647 | A-21448 | Thermo Fisher | 1:500 |
| Chromeo 494 Goat anti-mouse IgG | 15032 | ActiveMotif | 1:300 |
